# Towards a global perspective for *Salvia* L: Phylogeny, diversification, and floral evolution

**DOI:** 10.1101/2021.12.16.473009

**Authors:** Fatemeh Moein, Ziba Jamzad, Mohammadreza Rahiminejad, Jacob B. Landis, Mansour Mirtadzadini, Douglas E. Soltis, Pamela S. Soltis

## Abstract

**Premise of this study:** *Salvia* is the most species-rich genus in Lamiaceae, encompassing approximately 1000 species distributed all over the world. We sought a new evolutionary perspective for *Salvia* by employing macroevolutionary analyses to address the tempo and mode of diversification. To study the association of floral traits with speciation and extinction, we modeled and explored the evolution of corolla length and the lever-mechanism pollination system across our *Salvia* phylogeny.

**Methods:** We reconstructed a multigene phylogeny for 366 species of *Salvia* in the broad sense including all major recognized lineages and numerous species from Iran, a region previously overlooked in studies of the genus. Our phylogenetic data in combination with divergence time estimates were used to examine the evolution of corolla length, woody vs. herbaceous habit, and presence vs. absence of a lever mechanism. We investigated the timing and dependence of *Salvia* diversification related to corolla length evolution through a disparity test and BAMM analysis. A HiSSE model was used to evaluate the dependency of diversification on the lever-mechanism pollination system in *Salvia*.

**Key Results:** Based on recent investigations and classifications, *Salvia* is monophyletic and comprises ∼1000 species. Our inclusion, for the first time, of a comprehensive sampling for Iranian species of *Salvia* provides higher phylogenetic resolution for southwestern Asian species than obtained in previous studies. A medium corolla length (15-18mm) was reconstructed as the ancestral state for *Salvia* with multiple shifts to shorter and longer corollas. Macroevolutionary model analyses indicate that corolla length disparity is high throughout *Salvia* evolution, significantly different from expectations under a Brownian motion model during the last 28 million years of evolution. Our analyses show evidence of a higher diversification rate of corolla length for some Andean species of *Salvia* compared to other members of the genus. Based on our tests of diversification models, we reject the hypothesis of a direct effect of the lever mechanism on *Salvia* diversification.

**Conclusions:** Using a broader species sampling than previous studies, we obtained a well-resolved phylogeny for southwest Asian species of *Salvia*. Corolla length is an adaptive trait throughout the *Salvia* phylogeny with a higher rate of diversification in the South American clade. Our results suggest caution in considering the lever-mechanism pollination system as one of the main drivers of speciation in *Salvia*.

## 1. Introduction

Integrating molecular data with organismal traits can be used to address a major question in biology, “Is higher species diversity related to the presence of specific traits in that lineage?” (Pyron and Tubrin, 2014). Recently developed model-based approaches for estimating divergence times (BEAST: Drummond and Rambaut 2007; treePL: Smith and O’Meara 2010), diversification rates (MEDUSA: Alfaro et al., 2009; BAMM: Rabosky et al., 2014), and the effect of traits on diversification (FitzJohn et al., 2012; Beaulieu and O^’^Meara 2016; Caetano et al., 2018; Landis et al., 2018; Han et al., 2020) provide new opportunities to address this question. These methods have the advantage of providing estimates of the origin, divergence time, rate of diversification, and drivers of diversification among species.

There has been considerable recent interest in studying the association of floral traits and species richness in flowering plants (Vamosi et al., 2011; Van der Niet and Johnson 2014; Soltis & Soltis 2014; Saquet et al., 2017; Landis et al., 2018; Onstein 2019; Hernández and Wiens 2020). Interactions between flowers and their pollinators have spurred speciation and the evolution of novel floral variation (e.g., Stebbins 1970; Dodd et al., 1999; Crane et al., 1995; Crepet 2000; Soltis and Soltis 2004; Soltis et al., 2008; Ambruster 2014; Fenster et al., 2004; Smith 2010; Van der Niet and Johnson 2014). Some floral traits such as spur length, corolla shape, corolla length, and number of flowers are more often influenced by selection than other floral features (Yoshioka, 2007; Kacrowski et al., 2012; Landis et al., 2016). Floral specialization could potentially promote diversification by the evolution of adaptive floral traits through the establishment of reproductive isolation (Kay and Sargent 2009, Armbuster 2014; Serrano-Serrano et al., 2015). Several studies have also shown a correlation between flower specialization and rate of diversification (Fernández-Mazuecos et al., 2013; Ogutcen et al., 2014; Lagomarsino et al., 2016). For example, in genera of Neotropical Gesneriaceae including *Codonanthopsis* Mansf, *Codonanthe* (Mart.) Hanst, and *Nematanthus* Schard, species with hummingbird pollination syndromes have higher rates of diversification than close relatives pollinated by insects (Serrano-Serrano et al., 2015).

Lamiaceae (the mints) are the sixth largest family of flowering plants with over 7000 species distributed worldwide (Harley et al., 2004). Recently, Li et al. (2016, 2017) subdivided Lamiaceae into ten subfamilies and four unplaced genera based on a large-scale, plastid-based phylogenetic analysis, and this topology was largely corroborated by analysis of nuclear transcriptomes (Mint Evolutionary Genomics Consortium 2018). Within Lamiaceae, Nepetoideae is the largest subfamily with 105 genera and 3600 species, including well-known genera such as *Thymus* L. (thyme), *Ocimum* L. (basil), *Nepeta* L. (catnip), *Salvia* L. (sage), and *Lavandula* L. (lavender) (Harley et al., 2004).

*Salvia*, the largest genus in Lamiaceae as currently defined, includes approximately 1000 species, more than half of which are distributed in North and South America (Alziar, 1988- 1993). Morphologically, *Salvia* is highly diverse, particularly regarding specialized floral traits such as corolla color, corolla and tube length, flower shape and stamen structure (Wester and Claßen-Bockhoff, 2007; Reith et al., 2007; Will and Claßen-Bockhoff, 2015). Traditionally, *Salvia* was separated from other genera in Lamiaceae by possessing two fertile stamens with an elongated connective tissue. More that 80% of *Salvia* species are characterized by a special pollination system referred to as a lever mechanism (Walker et al., 2004; Harely et al., 2004; Claßen-Bockhoff et al., 2004; Walker and Sytsma, 2007). transfer to the stigma. The lever mechanism has the advantage of promoting successful pollination. In addition, this approach is efficient in pollen allocation and does not allow the pollinator to collect all of the pollen in one visit (Claßen-Bockhoff et al., 2003; Reith et al., 2007; Celep et al., 2014). A staminal lever is an advantage in *Salvia* due to the precise placement of pollen on bees while they are accessing the restricted nectar (Claßen-Bockhoff et al., 2004; Zhang et al. (2011) showed that removing the lever arms in *Salvia cyclostegia* resulted in lower fruit and seed set. Previous morphological studies of *Salvia* pollinators and floral traits hypothesized that the lever mechanism might play a role as a key innovation in promoting adaptive radiations (Claßen-Bockhoff et al., 2004; Will and Claßen-Bockhoff 2014). Based on phylogenetic results, the lever mechanism evolved in parallel in the Eastern and Western Hemispheres (Walker and Sytsma, 2007).

Since the initial phylogenetic study on Menthineae (Wagstaff and Olmstead, 1995), several studies have been performed based on nuclear and plastid regions with increasing taxonomic sampling of *Salvia* species (Walker and Sytsma, 2007; Takano and Okado 2011; Will and Claßen-Bockhoff, 2014; Drew and Sytsma, 2012; Will et al., 2015; Will and Claßen-Bockhoff, 2017; Hu et al., 2018; Drew et al., 2017; Fragoso-Martinez et al., 2017; Kriebel et al., 2019; Wu et al., 2021). In the first molecular study of *Salvia* (based on *rbcL* and the *trnL-trnF* regions), Walker and Sytsma (2004) found that *Salvia* is not monophyletic and recognized three clades: clade I includes many species of *Salvia* from the Eastern Hemisphere along with a Western Hemisphere lineage (8 species from former sect. *Heterosphacea* and subgen. *Salviaostrum*), clade II comprises North and South American species and includes subgen. *Calosphace* Benth. and subgen. *Audbertia* Benth, and clade III comprises species from eastern North Africa and southwestern Asia. *Rosmarinu*s L. and *Perovskia* Karel. were placed as sisters to clade I, while *Dorystaechas* Boiss. & Heldr. ex Benth, distributed in Turkey, was placed with clade II.

Walker and Sytsma (2007), with increased taxon sampling for *Salvia* and related genera in Menthineae, found that *Meriandra* Benth and *Dorystaechas* formed the sister clade to North and South American species of *Salvia* (clade II). They referred to *Zhumeria* Rech.f & Wendelbo (a monotypic genus endemic to Iran) along with southwest and East Asian *Salvia* as clade III. Will and Claßen-Bockhoff (2014) excluded the East Asian *Salvia* species from Walker and Sytsma’s (2007) clade III and considered them to represent an independent lineage (Clade IV). Will and Claßen-Bockhoff (2017) suggested breaking the large *Salvia* group into six genera: *Salvia sensu stricto*, *Ramonia* Raf., *Lasemia* Raf., *Glutinaria* Raf., *Pleudia,* and *Polakia*. However, they did not provide a taxonomic revision. Drew et al. (2017) embedded these five genera into a broadly defined *Salvia* and treated each as a subgenus. In recent phylogenetic studies of *Salvia*, Hu et al. (2018) and Kriebel et al. (2019) followed and updated the Drew et al. (2017) classification of *Salvia*, recognizing 11 subgenera. In this study, to maintain stability in taxonomic definition and nomenclature, we follow the broad definition of *Salvia* (Drew et al., 2017; Hu et al., 2018; Kriebel et al., 2019). A schematic diagram of changes in *Salvia* delimitation based on previous phylogenetic studies is provided in **Figure 1**.

**Fig 1:**
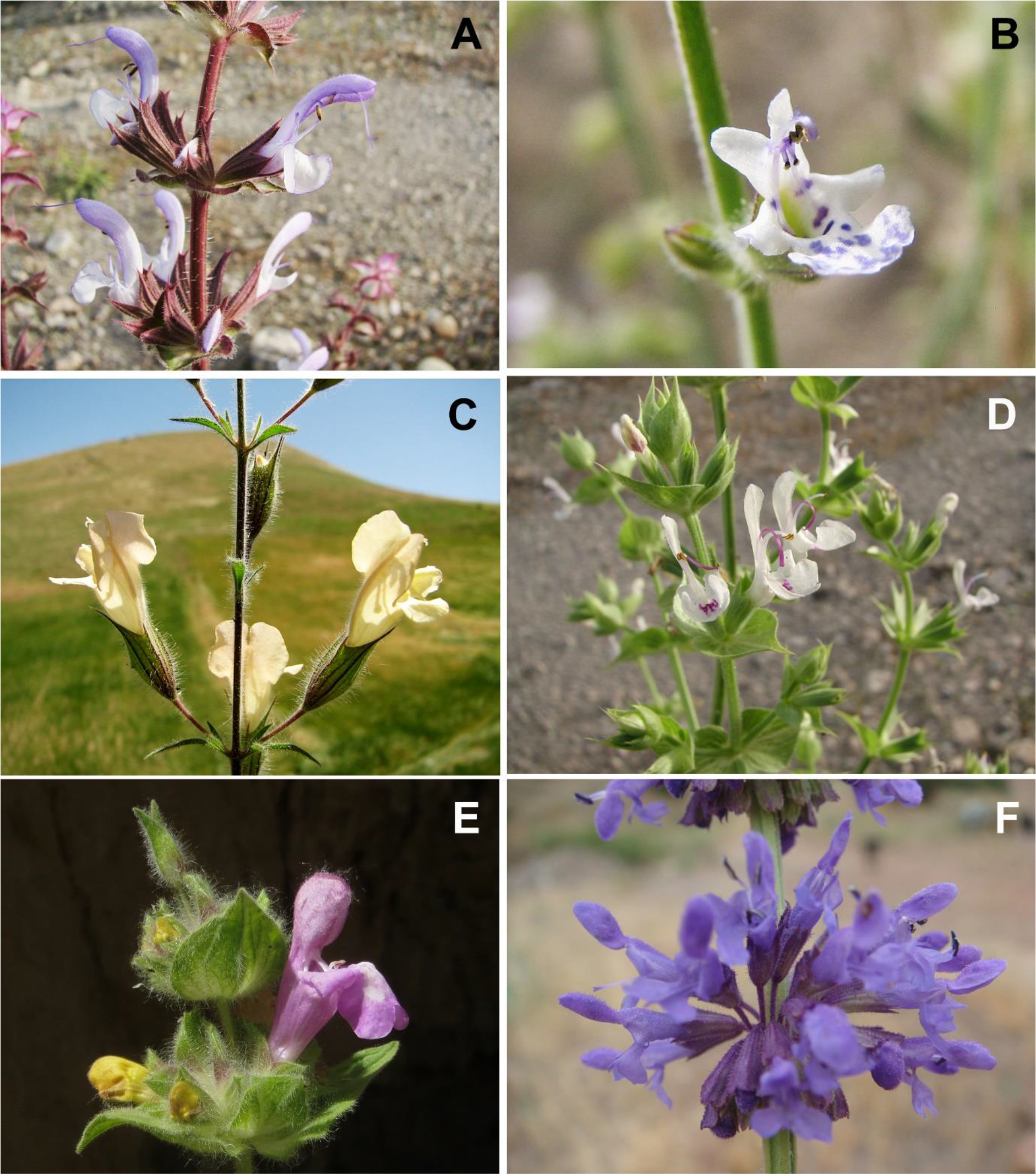
Phenotypic diversity in Iranian *Salvia*. A: *Salvia sclarea* (clade I), B: *Salvia aegyptiaca* (clade III) C: *Salvia aristata* (clade III), D*: Salvia macrosiphon* (clade I), E: *Salvia bracteata* (clade I), F: *Salvia verticillata* (clade I). A-E: Photos by M. Mirtajzadini, F: Photo by K. Safikhani.

**Fig. 2.**
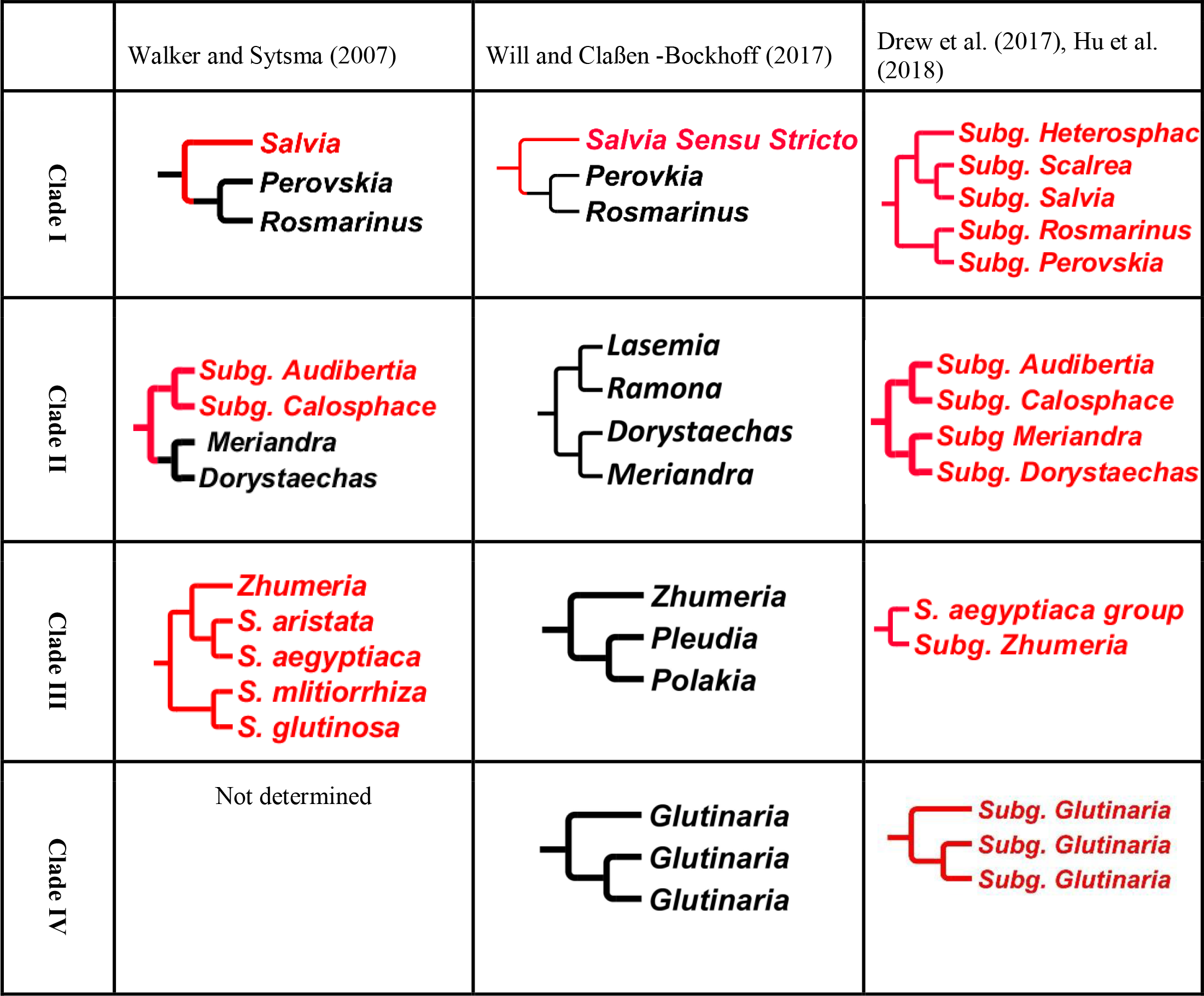
Schematic trees provide a summary of changes in *Salvia* delimitation based on previous phylogenetic studies (Walker and Sytsma, 2007; Will and Bockhoff, 2017; Drew et al., 2017). Those species that are classified under *Salvia* infrageneric delimitations are shown in red. Distinct genera from *Salvia* are indicated in black. Walker and Systma (2007) recognized three distinct clades for *Salvia* phylogeny embedded within five genera (*Perovskia, Rosmarinus, Dorystaechas, Meriandra* and *Zhumeria*). Will and Bockhoff (2017) identified just part of clade I as *Salvia* sensu stricto and split *Salvia* into six genera. Drew et al. (2017) maintained *Salvia* in the broad sense and treated the five genera in Walker and Sytsma (2007) as subgenera of *Salvia*. Hu et al. (2018) treated clade IV (from eastern Asia) as subg*. Glutinaria*.

Frequent endemism and enormous morphological diversity have made interpretation of the evolutionary patterns within *Salvia* challenging, particularly given the limited taxon sampling for some areas, such as southwestern Asia. To improve taxon sampling for southwestern Asia and to clarify patterns of morphological evolution and species diversification, we generated new sequences for 50 Iranian species of *Salvia* and reconstructed a phylogeny for 351 species overall. Notably, other recent phylogenetic analyses of *Salvia* differ in scope and emphasis from our investigation. Kriebel et al. (2019) studied the effect of biome shifts and pollinators on the radiation of *Salvia*. They found that shifts in pollination system are not correlated with species diversification, except in subgen. *Calosphac*e in the Western Hemisphere where species are pollinated by hummingbirds. Kriebel et al. (2020) showed that the respective floral morphospaces of the Western and Eastern Hemisphere *Salvia* are different. They inferred that these differences in flower morphology are linked with shifts from bee to bird pollination. In another recent study, Wester et al. (2020) found that shifts from bee- to bird-pollinated *Salvia* are mostly associated with floral structure rather than floral colors.

Despite valuable contributions, the relationship between the evolution of floral traits and patterns of *Salvia* diversification is not well understood. We used our new phylogenetic tree for *Salvia* to trace patterns of both character evolution and diversification. We primarily focus on the role of corolla length as one of the putative characters involved in *Salvia* diversification. This is also the first attempt to trace the evolutionary history of corolla length in *Salvia* and its association with diversification. Furthermore, we reconstructed the ancestral state for lever mechanism and habitat with greater taxon sampling than in previous work (Will and Claßen-Bockhoff 2014). In addition, we shed new light on the role of the pollination system in *Salvia* diversification. We statistically examine the longstanding hypothesis that the lever mechanism in *Salvia* flowers is correlated with high diversity and species richness.

## 2. Materials & Methods

### 2.1. Taxon Sampling

In total, 366 taxa representing 351 species covering all major areas of the geographic distribution of *Salvia* were used to reconstruct the phylogeny. As noted, we considered *Salvia* in the broad sense and included *Zhumeria*, *Meriandra*, *Rosmarinus*, and *Perovskia* (Drew et al., 2017, Kriebel et al., 2019). Following Drew and Sytsma (2012), we selected *Melissa* and *Lepechinia* as outgroups. We generated new sequences for many Iranian species of *Salvia*, including 50 species (59 accessions) for the external transcribed spacer (ETS) region of nuclear ribosomal DNA, 46 species (47 accessions) for ITS, and 35 species for the *ycf1-rps15* region of the plastome. The remaining sequences used here (representing 216 species) were obtained from GenBank. We concatenated all sequences for the three plastid regions (*rpl32*, *trnL-trnF*, *ycf1-rps15)* and two nuclear regions (ITS, ETS); the plastid and nuclear data sets were each analyzed separately and then combined, given the highly similar topologies obtained for each. That is, there was no strongly supported incongruence or conflict (hard incongruence sensu Seelanen et al., 1997) between nuclear and plastid trees. The newly generated sequences were deposited in GenBank. Corresponding information for each voucher specimen is provided in Table 1.

**Table 1:** Plant materials used in this study with their accession numbers. Newly generated sequences are shown in bold. TARI (National Herbarium of Iran, Research Institute of Forests and Rangelands)

|  | Voucher | ETS | ITS | Gen Bank<br>rpl32-trnl | trnl-trnf | Ycf1 |
| --- | --- | --- | --- | --- | --- | --- |
| *Lepechinia chamaedryoides* (Balb.) Epling (outgroup) | 1, 5) Drew and Sytsma 2011; 2) Walker and Sytsma 2007; 4) Walker et al., 2004. | JF301317 | DQ667231 | / | AY570459 | JF289031 |
| *Lepechinia leucopylloides* (Ramamoorthy, Hiriart & Medrano) B.T.Drew, Cacho & Sytsma, **comb. nov.** (outgroup) | 1, 2, 4, 5) Drew and Sytsma 2011; 3) Drew and Sytsma 2012. | JF301327.1 | JF301354 | JQ669348 | JF301390 | JF289047 |
| *Melissa officinalis* L. (outgroup) | 1, 2, 4, 5) Drew and Sytsma 2011; 3) Drew and Sytsma 2012. | JF301325 | JF301353 | JQ669335 | JF301386 | JF289042 |
| *S. africana-caerulea* L. | Will and Claßen-Bockhoff 2014. |  | KJ584204 | KJ7472171 | / | / |
| *Salvia absconditiflora* Greuter & Burdet (syn. *S. cryptantha* Montbret & Aucher ex Benth.) | Will and Claßen-Bockhoff 2017 | / | KU563839 | KU578211 | / | // |
| *Salvia acerifolia* B.L.Turner. | 2,4) Fragoso-Martinez et al., 2017. | / | MF664540 | / | MF663939 | / |
| *Salvia adenocaulon* H.P.Davis | Will and Claßen-Bockhoff 2017. | / | KU563828 | KU563828 | / | / |
| *Salvia adenophora* M.Martens & Galeotti | 2,4) Fragoso-Martinez et al., 2017. | / | MF622100 | / | MF663940 | / |
| *Salvia adenophylla* Hedge and Hub.-Mor. | Will and Claßen-Bockhoff 2017. | / | KU563789 | KU578218 | / | / |
| ***Salvia aegyptiaca* L.** | Iran: Bushehr, TARI (26835). | MK204892 | / | / | / | / |
| ***Salvia aegyptiaca*.** | Iran: Hormozgan, TARI (102853). | MK204891 | MK256969 | / | / | / |
| ***Salvia aethiopsis* L.** | Iran: TARI (6604). | MK204890 | / | / |  | MK240102 |
| *Salvia agnes* | 2,4) Fragoso-Martinez et al., 2017. |  | MF664541 |  | MF663941 |  |
| *Salvia akiensis* A.Takano, T.Sera et Kurosaki. | 1,2,4,5) Takano and Akiyama 2017. | LC060826 | LC060279 | / | LC124188 | LC060530 |
| *Salvia albicaulis* Benth. | 1,2,3) Will and Claßen-Bockhoff 2014. | KJ584257 | KJ584206 | KJ747274 | / | / |
| *Salvia albimaculata* Hedge and Hub.-Mor. | Will and Claßen-Bockhoff 2017. | / | KU563790 | KU578219 | / | / |
| *Salvia amethystina* Sm. | 2) Fragoso-Martinez et al., 2017. | / | MF664545 | / | / | / |
| *Salvia amplexicaulis* Lam. | Will and Claßen-Bockhoff 2017. | / | KU563829 | KU578151 | / | / |
| *Salvia anatolica* Hamzaog̃lu and Duran | Will and Claßen-Bockhoff 2017. | / | / | KU563840 | KU578221 | / |
| *Salvia apiana* Jeps. | 1,4,5) Walker et al., 2015; 2) Walker et al., 2004; 3) Will and Claßen-Bockhoff 2014. | KP852935 | DQ667214 | KJ747321 | KP852890 | KP853066 |
| *Salvia aramiensis* Rech.f. | Will and Claßen-Bockhoff 2017 | / | KU563791 | KU578247 | / | / |
| *Salvia arbuscular* Fernald | 2,4) Jenks et al., 2010 | / | HQ418846 | / | HQ418949 | / |
| *Salvia argentea* L. | Will and Claßen-Bockhoff 2014. | / | KJ584164 | KJ747299 | / | / |
| *Salvia aristate* Aucher ex Benth. | 1) Drew and Sytsma 2011; 2,4) Walker and Sytsma 2007; 5) Drew and Sytsma 2011 | JF301336 | D667280 | / | DQ667465 | JF289059 |
| ***Salvia aristata*** | Iran: Isfahan, TARI (12495). | MK204889 | / | / |  | MK240103 |
| *Salvia aspera* Fernald. | 2,4) Fragoso-Martinez et al., 2017. | / | MF664547 | / | MF663948 | / |
| *Salvia atrocyanea* Epling. | 2,4) Walker and Sytsma 2007. | / | DQ667270 | / | DQ667456 | / |
| ***Salvia atropatana* Bunge.** | Iran, TARI (88803) | MK204887 | MK213193 |  |  | / |
| ***Salvia atropatana*.** | Iran, TARI (29283) | MK204888 | / |  |  | / |
| *Salvia aucheri* Benth. | 2,4) Walker and Sytsma 2007; Will and Claßen-Bockhoff 2017 | DQ667286 | KU578248 | DQ667471 | / |  |
| *Salvia aurita* L. f. | Will and Claßen-Bockhoff 2014. | KJ584261 | KJ584218 | KJ747276 | / | / |
| *Salvia austriaca* Jacq. | 2,4) Walker and Sytsma 2007; 3) Will and Claßen-Bockhoff 2014. | / | DQ667323 | KJ747261 | DQ667512 | / |
| *Salvia axillaris* Moc. and Sessé ex Benth. | 1,3,5) Drew and Sytsma 2011. | JF301330 | MF664549 | JQ669366.<br>1 | MF663950 | JF301330 |
| *Salvia azurea* Michx. ex Vahl. | 2,4) Walker and Sytsma 2007. | / | DQ667317 | / | DQ667317 | / |
| *Salvia baimaensis* S.W.Su & Z.A.Shen | specimen_voucher="SBAI | / | JQ934103 | / | / | / |
| *Salvia ballotiflora* Benth. | 2,4) Jenks et al., 2010. | / | HQ418849 | / | HQ418952 | / |
| *Salvia bariensis* Thulin. | Will and Claßen-Bockhoff 2014. | KJ584262 | / | KJ747316 | / | / |
| ***Salvia bazmanica* Rech.f. & Esfand.** | Iran, TARI (43049). | MK204886 | / | / | / |  |
| *Salvia blepharochlaena* Bedge and Hub.-Mor. | Will and Claßen-Bockhoff 2017. | / | KU578210 | KU578210 | / | / |
| *Salvia blepharophylla* Hedge & Hub.-Mor. | 2) Jenks et al., 2010. | / | HQ418850 | / | HQ418953 | / |
| *Salvia brachyantha* | Will and Claßen-Bockhoff 2017 | / | KU563844 | KU578154 | / | / |
| ***Salvia brachysiphon* Stapf.** | Iran, TARI (3162). | MK204885 | MK213194 | / | / | MK240104 |
| ***Salvia brachysiphon*** | Iran, TARI (1145). | MK204884 | MK213195 | / | / | MK240105 |
| ***Salvia bracteata* Banks & Sol.** | Iran, TARI (16642). | MK204883 | / | / | / | MK240107 |
| *Salvia brandegeei* Munz. | 1,2,4,5) Walker and Sytsma 2005. | KP852949.1 | KP852783 | / | KP852896 | KP853067 |
| *Salvia breviflora* Moc. & Sessé ex Benth. | 2,4) Fragoso-Martinez et al., 2017. | / | MF664551 | / | MF663952 | / |
| *Salvia brevipes* Benth. | 2,4) Fragoso-Martinez et al., 2017. | / | MF664552 | / | MF663953 | / |
| *Salvia broussonetii* Benth. | Will and Claßen-Bockhoff 2014. | KJ584263 | KJ584225 | KJ747293 | / | / |
| *Salvia bucharica* M.Popov. | Will and Claßen-Bockhoff 2017. | / | KU563794 | KU578222 | / | / |
| *Salvia bulleyana* Diels. | Will and Claßen-Bockhoff 2017 | / | / | KU578203 | / | / |
| *Salvia cabulica* Benth. | Walker and Sytsma 2007. | / | DQ667287 | / | DQ667472 | / |
| *Salvia cacaliifolia* Epling | 2) Jenks et al., 2010; 4) Walker and Sytsma 2007. | / | HQ418851 | / | DQ667259 | / |
| *Salvia cadmica* Boiss. | Will and Claßen-Bockhoff 2017. | / | KU563795 | KU578223 | / | / |
| ***Salvia caespitosa* Montbret & Aucher.** | Iran, TARI (85247). | MK204882 | MK213197 | / | / | / |
| *Salvia californica* Brandegeei. | Drew and Sytsma 2015. | KP852951 | DQ667213 | / | DQ667424 | KP853068 |
| *Salvia candelabrum* Boiss. | Will and Claßen-Bockhoff 2014. | / | KJ584190 | KJ747255 | / | / |
| *Salvia candicans* M.Martens and Galeotti. | 2,4) Fragoso-Martinez et al., 2017. | / | MF664557 | / | MF663958 | / |
| *Slavia candidissima* Vahl. | Walker and Sytsma 2007 | / | DQ667261 | / | DQ667447 |  |
| *Salvia carduacea* Benth. | Walker et al., 2015. | / | KP852785 | / | KP852900 | KP853069 |
| *Salvia carnea* Kunth. | 2,4) Jenks et al., 2010. | / | HQ418854 | / | HQ418957 | / |
| *Salvia cassia* Sam. ex Rech.f. | Will and Claßen-Bockhoff 2017. | / | KU563845 | KU578190 | / | / |
| *Salvia castanea* Diels | 2) Will and Claßen-Bockhoff 2017; 4) Wang et al., 2013. | / | KU563781 | / | KC414279 | / |
| *Salvia cavaleriei* H.Lév. | 1,4) Wang et al., 2013. |  | KC473232 |  | KC414280 |  |
| *Salvia cedrosensis* Greene. | 1,5) Walker et al., 2015; 2) Walker and Sytsma 2007; 4) Murphy and Bola 2012. | KP852953 | DQ667228 | / | JQ888128 | KP853070 |
| ***Salvia ceratophylla* L.** | Iran, TARI, 17238B. | MK204881 | MK213198 | / | / | MK240107 |
| *Salvia chamaedryoides* Cav | 2) Jenks et al., 2010; 4) Walker et al., 2004 | / | HQ418855 | / | AY570471 | / |
| *Salvia chamelaeagnea* Berg. | Will and Claßen-Bockhoff 2014 | KJ584268 | KJ584210 | KJ747289 | / | / |
| *Salvia chienii* E. Peter. | Will and Claßen-Bockhoff 2014 | / | KJ584250 | KJ747322 |  |  |
| *Salvia chinensis* Benth. | PS0121MT04 | / | FJ883503 | / | / | / |
| *Salvia chionantha* Boiss. | Will and Claßen-Bockhoff 2017 | / | KU563846 | KU578155 | / | / |
| *Salvia chionoeplica* Epling. | 1,2,3) Will and Claßen-Bockhoff 2014; 4) Walker et al., 2004; 5) Walker et al., 2015. | KJ747318 | KJ584188 | KJ747318 | AY570472 | KP853071 |
| ***Salvia chloroleuca* Rech.f. & Aellen.** | Iran, TARI (36026). | MK204879 | MK212199 | / | / | MK240109 |
| ***Salvia chloroleuca*.** | Iran, TARI (12691). | MK204880 | / | / | / | MK240108 |
| ***Salvia chorassanica* Bunge.** | Iran, TARI (5354). | MK204878 | MK213200 | / | / | MK240110 |
| *Salvia chrysophylla* Stapf. | Will and Claßen-Bockhoff 2017. | / | KU563848 | KU578157 | / | / |
| *Salvia cinnabarina* M.Martens and Galeotti. | 2,4) Fragoso-Martinez et al., 2017. | / | MF664559 | / | MF663960 | / |
| *Salvia clevelandii* (A. Gray). | 1) JBW 3079; 2,4) Fragoso-Martinez et al., 2017. | KP853032 | MF664560 | / | MF663961 | / |
| *Salvia clinopodioides* Kunth. | 2,4) Fragoso-Martinez et al., 2017 | / | MF664561 | / | MF663962 | / |
| *Slavia coccinea* Buc'hoz ex Etl. | 2) Trusty et al., 2004; 4) Wang et al., 2013. | / | AY506651 | / | KC414281 | / |
| *Salvia columbariae* Benth. | 4) Walker et al., 2015 | KP852960.1 | KP852793 | / | KP852905 | KP853073 |
| ***Salvia compressa* Vent.** | Iran: Hormozgan, TARI (102856) | MK204877 | MK213201 | / | / | / |
| *Salvia concolor* Lamb. ex Benth. | 2,4) Jenks et al., 2010 | / | HQ418858 | / | HQ418961 | / |
| *Salvia confertiflora* Pohl. | 2,4) Jenks et al., 2010. | / | HQ418859 | / | HQ418962 | / |
| *Salvia confertispicata*. | 2,4) Fragoso-Martinez et al., 2017 | / | MF664563 | / | MF663964 | / |
| *Slavia congestiflora* Epling. | 2,4) Fragoso-Martinez et al., 2017. | / | MF664564 | / | MF663965 | / |
| *Salvia connivens* Epling. | 2,4) Fragoso-Martinez et al., 2017 | / | MF664565 | / | MF663966 | / |
| *Salvia corrugata* Vahl. | 2) Fragoso-Martinez et al., 2017; 4) Walker et al., 2004. |  | MF622122 |  | AY570476 |  |
| *Salvia cuspidata* (Benth.) J.R.I.Wood. | 2,4) Fragoso-Martinez et al., 2017. | / | MF664568 | / | MF663970 | / |
| *Salvia cyanescens* Boiss. and Bal. | Will and Claßen-Bockhoff 2017. | / | KU563849 | KU578158 | / | / |
| *Salvia cyclostegia* E. Peter. | 2,4) Wang et al., 2013. | / | KC473274 | / | KC414282 |  |
| *Salvia cynica* Dunn. | Walker and Sytsma 2007. | / | DQ667332 | / | DQ667521 |  |
| *Salvia dabieshanesis* J.Q.He. | PS1723MT01 |  | FJ883505 | / | / | / |
| *Salvia daghestanica* Sosn. | Walker and Sytsma 2007. | / | KJ584187 | KJ747308 | DQ667444 | / |
| *Salvia densiflora* Benth | 2,4) Fragoso-Martinez et al., 2017. | / | MF664570 | / | MF663972 | / |
| *Salvia deserta* omit. | Will and Claßen-Bockhoff 2017. | / | KJ584176 | KJ747263 |  |  |
| *Salvia deserti* Decne. | 2,3) Will and Claßen-Bockhoff 2014. | KJ584270 | / | KJ747312 | / | / |
| *Salvia dianthera* Roth ex Room, and Schult. | 2, 4) Walker and Sytsma 2007; 5) Drew and Sytsma 2011. | JF301326.1 | DQ667329 | / | DQ667518 | JF289044 |
| *Salvia dichroantha* Stapf. | Will and Claßen-Bockhoff 2017 | / | KU578159 | KU56830 |  |  |
| *Salvia digitaloides* Diels. | 2) Walker & Sytsma 2007; 4) Walker & Sytsma 2004. | / | / | DQ667255 | AY570477 | / |
| *Salvia discolor* Sessé & Moc. | 2,4) Jenks et al., 2010. | / | HQ418860 | / | HQ418963 | / |
| *Salvia disermas* L. | 1,2,3) Will and Claßen-Bockhoff 2014; 4) Walker et al., 2004. | KJ584271 | KJ584179 | KJ747296 | AY570478 | / |
| *Salvia disjuncta* Fernald. | 2,3) Will and Claßen-Bockhoff 2017; 4) Fragoso-Martinez et al., 2017. | / | KU563882 | KU578197 | MF663975 | / |
| *Salvia divaricata* Montbret & Auch. ex Benth. | Will and Claßen-Bockhoff 2017 | / | / | KU578226 | / | / |
| *Salvia divinorum* Epling & Játiva. | 2,4) JBW 2330 | / | HQ418861 | / | DQ667440 | / |
| *Salvia dolomitica* Codd. | 1,3) Will and Claßen-Bockhoff 2014; 2,4) Walker and Sytsma 2007; | KJ584274 | DQ667322 | KJ747290 | DQ667511 | / |
| *Salvia dombeyi* Epling. | 2,4) Jenks et al., 2010 | / | HQ418862 | / | HQ418965 | / |
| *Salvia dorisiana* Standl. | 2,4) Jenks et al., 2010 | / | HQ418863 | / | HQ418966 | / |
| *Salvia dorrii* (Kellogg.) Abrams. | 1,4,5) Walker et al., 2015; 2) Walker and Sytsma 2007 | KP853037 | DQ667229 | / | KP852907 | KP853074 |
| *Salvia dorystaechas* B. T. Drew | 2) Will and Claßen-Bockhoff 2014; 3) Drew and Sytsma 2012; 4) Walker et al., 2004; 5) Drew and Sytsma 2011 | KJ584257 | KJ584284 | JQ669302 | AY570454 | JF289014 |
| ***Salvia dracocephaloides* Boiss.** | Iran, TARI (30192). | MK204876 | Mk213202 | / | / | MK240111 |
| *Salvia elegans* Vahl. | 2,4) Fragoso-Martinez et al., 2017. | / | MF622127 | / | MF663978 | / |
| *Salvia engelmannii* A. Grey. | Will and Claßen-Bockhoff 2017 | / | KU563870 | Ku578163 | / | / |
| ***Salvia eremophila* Boiss.** | Iran, TARI (41741). | MK204875 | / | / | / | MK240112 |
| *Salvia eremostachya* Jeps. | 1,4,5) Walker et al., 2015; 2) Walker and Sytsma 2007. | KP853039 | DQ667232 | / | KP852910 | KP853075 |
| *Salvia eriophora* Boiss. and Kotschy. | Will and Claßen-Bockhoff 2017 | / | KU563850 | KU578167 |  |  |
| *Salvia erythrostoma* Rusby. | Jenks et al., 2010. | / | HQ418864 | / | HQ418967 | / |
| *Salvia euphratica* Montbret and Aucher ex Benth. | Dizkirici et al., (2015). | / | KM519756 | KU578227 | KM519770 | / |
| *Salvia evansiana* Hand.-Mazz. | 2,3) Will and Claßen-Bockhoff 2014; 3) Zhong et al., 2010. | / | KJ584251 | KJ747323 | FJ593462.1 | / |
| *Salvia exserta* Grieseb. | 2,4) Fragoso-Martinez et al., 2017 | / | MF622128 | / | MF663979 | / |
| *Salvia farinacea* Benth. | 2) Chen et al., 2010; 4) Walker et al., 2004. | / | FJ546871 | / | AY570479 | / |
| *Salvia filifolia* Ramamoorthy. | 2,4) Fragoso-Martinez et al., 2017. | / | MF664576 | / | MF663980 | / |
| *Salvia flocculosa* Benth. | 2,4) Fragoso-Martinez et al., 2017. | / | MF664578 | / | MF663982 | / |
| *Salvia formosa* L'Hér. | 2) Fragazo –Martinez et al., 2017. | / | MF622131 | / | / | / |
| *Salvia frigida* Boiss. | Will and Claßen-Bockhoff 2017. | / | KU563851 | KU578168 | / | / |
| *Salvia fruticosa* Miller. | Will and Claßen-Bockhoff 2014. | / | KJ584195 | KJ747256 | / |  |
| *Salvia fulgens* Cav. | 2) Walker and Sytsma 2007; 4) Benitez-Vieyra et al., 2014. | / | MF622133 | / | KJ473988 | / |
| *Salvia funerea* M. E. Jones. | Walker et al., 2015. | KP853041 | KP852812 | / | KP852911 | KP853076 |
| *Salvia galloana* B.L.Turner. | 2,4) Fragoso-Martinez et al., 2017. | / | MF664581 | / | MF663985 | / |
| *Salvia garipensis* E.Meyer ex Benth | Walker and Sytsma 2007. | / | DQ667281 | / | DQ667466 | / |
| *Salvia geminata* Thulin. | 1) Will and Claßen-Bockhoff 2014. | KJ584276 | / | / | / | / |
| *Salvia gesneriiflora* Lindl. & Paxton. | 2,4) Fragoso-Martinez et al., 2017. | / | MF622133 | / | MF663986 | / |
| *Salvia glabrescens* var. *repens* (Koidz.) Kurosaki. | Sudarmono and Okada 2007. | LC060829.1 | AB295104 | / | AB295089 | LC060533 |
| ***Salvia glutinosa* L.** | Iran, TARI (21565). | MK204873 | MK213203 | / | / | Mk240113 |
| *Salvia glutinosa*. | 2) Will and Claßen-Bockhoff 2014; 4) Walker et al., 2004; 5) Drew and Sytsma 2011. | / | KU563774 | / | AY570480 | JF289061 |
| *Salvia gracilliramulosa* Epling & Játiva. | 2.4) Walker and Sytsma 2007. | / | DQ667276 | / | DQ667461 | / |
| *Salvia gravida* A.Gray. | 2,4) Jenks et al., 2010. | / | HQ418868 | / | HQ418971 | / |
| *Salvia greatae* Brandegees. | 1,5) Drew and Sytsma 2011; 2) Walker and Sytsma 2007; 3) Drew and Sytsma 2012; 4) Walker et al., 2004. | JF301331 | DQ667215 | JF289062 | AY570481 | JF289062 |
| *Salvia greggii* A.Gray. | 2,4) Jenks et al., 2010. | / | HQ418870 | / | HQ418972 | / |
| *Salvia greuiifolia* S. Moore. | 2,4) Jenks et al., 2010. | / | HQ418871 | / | HQ418973 | / |
| ***Salvia grossheimii* Sosn.** | Iran, TARI (84031). | MK204832 | MK213204 | / | / | / |
| *Salvia guadalajarensis* Briq. | 2,4) Fragoso-Martinez et al., 2017. | / | MF664584 | / | MF663989 | / |
| *Salvia guaranitica* Briq. | 2,4) Wang et al., 2013. | / | KC473237 | / | KC414285 |  |
| *Salvia handelii* E.Peter. | SHAN |  | JQ934124 | / | / | / |
| *Salvia hayatana* Makino ex Hayata. | 2,4) Sudarmono and Okada 2007; 5) Takano 2017. | / | AB295099 | / | AB295084 | / |
| *Salvia heerii* Regel. | 2,4) Fragoso-Martinez et al., 2017. | / | MF664587 | / |  | / |
| *Salvia heldreichiana* Boiss. ex Benth | Will and Claßen-Bockhoff 2017. | / | KU563799 | KU578246 | / | / |
| *Salvia henryi* Gray. | Will and Claßen-Bockhoff 2017. | / | KU563875 | KU578165 | / | / |
| *Salvia herbacea* Benth. | 2,4) Fragoso-Martinez et al., 2017. | / | MF664589 | / | MF663994 | / |
| *Salvia herbanica* A.Santos and M.Ferna'ndez. | Will and Claßen-Bockhoff 2014. | KJ584278 | KJ584246 | KJ747313 | / | / |
| *Salvia heterochroa* E. Peter. | 2,3) Will and Claßen-Bockhoff 2014. | / | KJ584252 | KJ747324 | / | / |
| *Salvia heterochroa* E.Peter. | Will and Claßen-Bockhoff 2017 | / | KJ584252 | KJ747324 | / | / |
| *Salvia heterofolia* Epling & Mathias. | 2,4) Fragoso-Martinez et al., 2017. | / | MF664590 | / | MF663995 | / |
| *Salvia hians* Royle ex Benth. | 4) Walker et al., 2004. | / | DQ763239 | / | AY570483 | / |
| *Salvia hidalgensis* Miranda. | 2,4) Fragoso-Martinez et al., 2017. | / | MF664591 | / | MF663996 | / |
| *Salvia hintonii* Epling. | 2,4) Fragoso-Martinez et al., 2017. | / | MF664592 | / | MF663997 | / |
| *Salvia hirtella* Vahl | 2,4) Walker and Sytsma 2007. | / | DQ667326 | / | DQ667515 | / |
| *Salvia hispanica* L. | 2,4) Fragoso-Martinez et al., 2017. | / | MF664593 | / | MF663998 | / |
| *Salvia honania* L.H.Bailey. | PS1722MT01 | / | FJ883513 | / | / | / |
| *Salvia huberi* Hedge. | Will and Claßen-Bockhoff 2017. | / | KU563800 | KU578228 | / | / |
| *Salvia hylocharis* Diels | Wang et al., 2013. | / | KC414286 | / | KC473238 | / |
| *Salvia hypargeia* Fisch. and Mey. | Will and Claßen-Bockhoff 2017. | / | KU563876 | KU578196 | / | / |
| ***Salvia hypochionaea* Boiss.** | Iran, TARI (30437). | MK204870 | MK213206 | / | / | / |
| ***Salvia hypoleuca* Benth.** | Iran, TARI (54151). | MK204869 | MK213207 | / | / | MK240115 |
| ***Salvia indica* L.** | Iran, TARI (90002). | MK204868 | MK213208 | / | / | MK240116 |
| *Salvia interrupta* Schousb. | Will and Claßen-Bockhoff 2014. | / | KJ584191 | KJ747265 | / |  |
| *Salvia isensis* Nakai ex Hara. | Takano and Akiyama 2017. | LC060831.1 | LC060730 | / | LC124190 | LC060535 |
| ***Salvia jamzadii* Mozaff.** | Iran, TARI (61992). | MK204867 | MK213209 |  |  | MK240117 |
| *Salvia japonica* f. longipes (Nakai) Sugimoto. | 1,4,5) Sudarmono and Okada (2007); 2) Takano and Akiyama 2017. | LC060835.1 | AB266239 | / | LC124191 | LC060537 |
| *Salvia judaica* Boiss. | Will and Claßen-Bockhoff 2017. | / | KJ584241 | KU578160 | / | / |
| *Salvia jurisicii* Košanin. | Will and Claßen-Bockhoff 2017 | / | KU563831 | KU578173 | / | / |
| *Salvia karwinskii* Benth. | 2,4) Fragoso-Martinez et al., 2017 | / | MF622144 | / | MF664003 | / |
| *Salvia kiaometiensis* H. Lév. | 2,4) Wang et al., 2013 | / | KC473239 | / | KC414287 | / |
| *Salvia Koyamae* Makino. | 2,4) Takano and Okada 2011 |  | AB541114 | AB541142 | / | LC060540 |
| *Salvia kronenburgii* Rech. f. | Will and Claßen-Bockhoff 2017 | / | KM519759 | KU578245 | KM519773 | / |
| *Salvia kurdica* Boiss. and Hohen ex Benth. S. | Will and Claßen-Bockhoff 2017 | / | KU563821 | KU578212 | / | / |
| *Slavia lasiantha* Benth. | 2,4) Walker and Sytsma 2007. | / | DQ667300 | / | DQ667486 | / |
| ***Salvia lachnocalyx* Hedge.** | Iran, TARI (83023). | MK204866 | MK213210 |  |  | MK240118 |
| *Salvia lachnostachys* Benth. | 2,4) Fragoso-Martinez et al., 2017 | / | MF664598 | / | MF664005 | / |
| *Salvia lanceolata* Lam. | 2,3) Will and Claßen-Bockhoff 2014 | / | KJ584201 | KJ747277 | / | / |
| *Salvia leptostachys* Benth. | 2,4) Fragoso-Martinez et al., 2017 | / | MF664603 | / | MF664010 | / |
| ***Salvia leriifolia* Benth.** | Iran, TARI (35583). | MK204865 | MK213211 |  |  | / |
| *Salvia leucantha* Cav. | 2,4) Jenks et al., 2010. | / | HQ418875 | / | HQ418977 | / |
| *Salvia leucodermis* Baker. | Will and Claßen-Bockhoff 2014. | KJ747280 | KJ584220 | KJ747280 | / | / |
| *Salvia leucophylla* Greene. | 2) Walker and Sytsma 2007; 4) Walker et al., 2004 | / | DQ667210 | / | KP852913 | KP853077 |
| ***Salvia limbata* C.A.Mey.** | Iran, TARI (27761). | MK204862 | MK213212 | / | / | MK240120 |
| ***Salvia limbata*** | Iran, TARI (30364). | MK204864 | MK213213 |  |  | Mk24011 |
| ***Salvia limbata*** | Iran, TARI (85267). | MK204863 | / | / | / | / |
| *Salvia littae* Vis. | 2,4) Fragoso-Martinez et al., 2017. | / | MF622150 | / | MF664014 | / |
| *Salvia longispicata* M.Martens & Galeotti. | 2,4) Jenks et al., 2010. | / | HQ418876 | / | HQ418978 | / |
| *Salvia lophanthoides* Fernald. | 2,4) Fragoso-Martinez et al., 2017. | / | MF664607 | / | MF664017 | / |
| *Salvia lutea* L. | Will and Claßen-Bockhoff 2014. | KJ747273 | KJ584205 | KJ747273 | / | / |
| *Salvia lutescens* var. *lutescens* Koidz. | Takano and Akiyam 2017; 2,4) Sudarmono and Okada 2007. | LC060845.1 | AB266232 | / | AB266232 | / |
| *Salvia lyrata* L. | Will and Claßen-Bockhoff 2017. | / | KU563873.1 | KU578166 | / | / |
| ***Salvia macilenta* Boiss.** | Iran, TARI (102851). | MK204861 | MK213214 | / | / | MK240122 |
| ***Salvia macrochlamys* Boiss. & Kotschy** | Iran, TARI (102852) | MK204860 | MK213215 | / | / | MK240123 |
| *Salvia macrophylla* Bent. | 2,4) Jenks et al., 2010. | / | HQ418877 | / | HQ418979 |  |
| ***Salvia macrosiphon* Boiss.** | Iran, TARI (58399). | MK204859 | MK213216 | / | / | MK240124 |
| *Salvia madrensis* Seem. | 2,4) Jenks et al., 2010. | / | HQ418878 | / | HQ418980 | / |
| ***Salvia majdae* (Rech.f. & Wendelbo) Sytsma.** | Iran: Hormozgan, Geno mountain, Mirtajzadin 201. | MK204858 | MK256967 | / | / | / |
| *Salvia marashica* Ilçim, Celep and Dogan. | Will and Claßen-Bockhoff 2017. | / | KU563802 | KU578230 | / | / |
| *Salvia margaritae* Botsch. | 2,3) Will and Claßen-Bockhoff 2017. | KU563880 | / | KU578201 | / | / |
| *Salvia maximowicziana* Hemsl. | 1,3,4) Deng et al., 2015. | KM886617 | PS1730MT01 | KM886851 | KM886650 |  |
| *Salvia maymanica* Hedge. | Will and Claßen-Bockhoff 2017. | / | KU563805 | KU578231 | / | / |
| *Salvia meilienis* S.W.Su. | PS1719MT01 | / | FJ546867 | / | / | / |
| *Salvia mellifera* E. Greene. | 1,4) Walker et al., 2015; 2) Jenks et al., 2011; 3) Drew and Sytsma 2012; 5) Drew et al., 2011 | KP852989 | HQ418879 | JQ669368 | KP852916 | JF289064 |
| *Salvia merjamie* Forsk. | Will and Claßen-Bockhoff 2014. | KJ584286 | KJ584184 | KJ747297 | / | / |
| *Salvia mexicana* L. | 2,4) I. Fragoso-Martinez 79 (FCME). | / | MF664611 | / | MF664021 | / |
| *Salvia microphylla* Kunth | 2,4) Fragoso-Martinez et al., 2017. | / | MF664022 | / | MF663986 | / |
| ***Salvia microstegia* Boiss. & Balansa** | Iran, TARI (5397). | MK204857 | MK213216 | / | / | / |
| *Salvia minarum* Briq. | 2,4) Fragoso-Martinez et al., 2017 | / | MF664613 | / | MF664023 | / |
| ***Salvia mirzayanii* Rech.f. & Esfand.** | Iran, TARI (41724). | MK204856 | MK213217 | / | / | / |
| *Salvia misella* Kunth. | 2,4) Fragoso-Martinez et al., 2017 | / | MF664614 | / | / | / |
| *Salvia mocinoi* Benth. | 2) Fragoso-Martinez et al., 2017; 4) Walker and Sytsma 2007 | / | MF664615 | / | DQ667459 | / |
| *Salvia modesta* Boiss. | Will and Claßen-Bockhoff 2017 | / | KU563856 | KU578175 | / | / |
| *Salvia mohavensis* E. Greene. | 2) Walker and Sytsma 2007; 1,4,5) Walker et al., 2015 | KP852997 | DQ667212 | / | KP852920 | KP853078 |
| *Salvia moniliformis* Fernald. | 2,3) Will and Claßen-Bockhoff 2017 | / | KU563884 | KU578198 | / | / |
| *Salvia montbretii* Benth. | Will and Claßen-Bockhoff 2017 | / | KU563869 | KU578195 | / | / |
| *Salvia muirii* L.Bolus. | Will and Claßen-Bockhoff 2014. | KJ584287 | KL584208 | KJ747283 | / | / |
| ***Salvia multicaulis* Vahl.** | Iran, TARI (102845) | / | MK213218 | / | / | / |
| *Salvia munzii* Epling. | 2) Walker and Sytsma 2007; 1,5) Walker et al., 2015; 4) Fragoso-Martinez et al., 2017. | KP853000 | DQ667224 | / | MF664026 | KP853079 |
| *Salvia namaensis* Schinz. | Will and Claßen-Bockhoff 2014. | KJ584289 | KJ584200 | KJ747284 | / | / |
| *Salvia nana* Kunth. | 2,4) Fragoso-Martinez et al., 2017 | / | MF664618 | / | MF664029 | / |
| ***Salvia nemorosa* L.** | Iran, TARI (43572) | MK204855 | MK213219 | / | / | MK240125 |
| *Salvia nervosa* Benth. | 2,4) Fragoso-Martinez et al., 2017. | / | MF664619 | / | MF664031 | / |
| *Salvia nilotica* Juss. ex Jacq. | 1,2,3) Will and Claßen-Bockhoff 2014; 4) Walker and Sytsma 2004 | KJ747281 | KJ584229 | KJ747258 | AY570487 | / |
| *Salvia nipponica* Miq. var. kisoensis | 1,5) Takano and Akiyam 2017; 2) Takano and Okada 2011 | LC060848 | AB295101 | / | / | LC060552 |
| *Salvia nitida* (M.Martens & Galeotti) Benth. | 2,4) Fragoso-Martinez et al., 2017 | / | MF664620 | / | MF664032 | / |
| *Salvia nubicola* Wall. ex Sweet | 2,4) Will and Claßen-Bockhoff 2017 |  | KU563786 | KU578205 | / | / |
| *Salvia nutans* L. | Will and Claßen-Bockhoff 2017 | / | KU563832 | KU578176 | / | / |
| *Salvia nydeggeri* Hub.-Mor. | Will and Claßen-Bockhoff 2017 | / | KU563803 | KU578233 | / | / |
| *Salvia oaxacana* Fernald. | 3) Will and Bockhoff 2017. | / | / | KU578199 | HQ418983 | / |
| *Salvia occidentalis* Sw. | 2,4) Jenks et al., 2010. | / | HQ418882 | / | HQ418984 | / |
| *Salvia officinalis* L. | 1, 4, 5) Drew and Sytsma 2011; 2) Walker and Sytsma 2007 | JF301332 | DQ667225 |  | JF301398 | JF289065 |
| ***Salvia oligophylla* Aucher ex Benth.** | Iran: Zanjan, Mirtajzadini | MK204853 | MK256968 |  |  | MK240126 |
| *Salvia omeiana* E. Peter. | Voucher: SOME | / | JQ934139 | EU200176 | / | / |
| *Salvia omerocalyx* Hayata var omerocalyx | 1,5) Takano 2017; 2,4) Takano and Okada 2011 | LC060852.1 | AB353205 | / | AB353195 | LC060557 |
| *Salvia oppositiflora* Ruiz & Pav. | 2,4) Jenks et al., 2010 | / | HQ418883 | / | HQ418985 | / |
| *Salvia orbignaei* Benth. | 2) Fragoso-Martinez et al., 2017; 4) Walker and Sytsma 2007 | / | DQ667279 | / | MF664033 | / |
| *Salvia ovalifolia* A.St.-Hil. ex Benth. | 4) Walker and Sytsma 2007 | / | DQ667315 | / | DQ667502 | / |
| *Salvia oxyphora* Briq. | 2,4) Jenks et al., 2010 | / | HQ418884 | / | HQ418986 | / |
| *Salvia pachyphylla* Munz. | 1,5) Walker et al., 2015; 2) Walker and Sytsma 2007 | KP853004 | / | / | DQ667431 | KP853080 |
| *Salvia palaestina* Benth. | Will and Claßen-Bockhoff 2014 | / | KJ584175 | KJ74304 | / | / |
| *Salvia pallida* Benth. | 2,4) Fragoso-Martinez et al., 2017 | / | MF664623 | / | MF664035 | / |
| *Salvia paohsingensis* C.Y.Wu. | 2,4) Wang et al., 2013 | / | KC473252 | / | KC414297 | / |
| *Salvia patens* Cav. | 1,5) Drew and Sytsma 2011; 2) Jenks et al. 2010; 3) Drew and Sytsma 2012; 4) Walker and Sytsma 2007 | JF301333 | HQ418885 | JQ669370 | DQ667442 | JF289066 |
| *Salvia pauciflora* Kunth. | 2,4) Wang et al., 2013. | / | KC473254 | / | KC414299 | / |
| *Salviapentstemonoides* K.Koch and C.D.Bouché. | 3) Will and Claßen-Bockhoff 2014; 2,4) Walker and Sytsma 2007. | / | DQ667221 | KU578162 | AY570489 | / |
| *Salvia perlonga* Fernald. | 2,4) Fragoso-Martinez et al., 2017. | / | MF664627 | / | MF664040 | / |
| *Salvia personata* Epling. | 2,4) Walker and Sytsma 2007. | / | DQ667269 | / | DQ667455 | / |
| ***Salvia persopolitana* Boiss.** | Iran, TARI (102854). | MK204852 | MK213220 |  |  | MK240127 |
| *Salvia phlomoides* Asso. | Will and Bockhof 2014. | / | KJ584186 | KJ747309 | / | / |
| *Salvia pinnata* L. | Will and Claßen-Bockhoff 2017. | / | KU563798 | KU578217 | / | / |
| *Salvia platystoma* | 2,4) Walker and Sytsma 2007. | / | DQ667277 | / | DQ667462 | / |
| *Salvia plebeia* R.Br. | 1,5) Takano 2017; 2) Will and Claßen-Bockhoff 2017; 3) Wang et al., 2013. | LC060859.1 | KU563788 | / | KC414300 | LC060563 |
| *Salvia plectranthoides* Griff. | 2,3) Will and Claßen-Bockhoff 2017. | / | KU563787 | KU578207 | / | / |
| ***Salvia poculata* Nábelek.** | Iran, TARI (102848). | MK204851 | MK213221 | / | / | / |
| *Salvia polystachya* Cav. | 1,5) Drew and Sytsma 2011; 2,4) Fragoso-Martinez et al., 2017; 3) Drew and Sytsma 2012. | JF301334 | MF664631 | JQ669371 | MF664044 | JF289067 |
| *Salvia potentillifolia* Boiss. and Heldr. ex Benth. | Will and Claßen-Bockhoff 2017. | / | KU563807 | KU578235 | / | / |
| *Salvia pratensis* L. | Will and Claßen-Bockhoff 2017. | / | KU563835 | KU578180 | / | / |
| *Salvia prattii* Hemsl. | 2,3) Will and Claßen-Bockhoff 2017. | / | KU578206 | / | KU563784 | / |
| *Salvia priontis* Hance. | specimen_voucher="PS1711MT01 | / | FJ883527 | / | / | / |
| *Salvia procurrens* Benth. | 2,4) Fragoso-Martinez et al., 2017. | / | MF664633 | / | MF664046 | / |
| *Salvia prunelloides* Kunth. | 2) Walker and Sytsma 2007; 4) Fragoso-Martinez et al., 2017. | / | DQ667275 | / | MF664048 | / |
| *Salvia przewalskii* Maxim. | 1,5) Drew and Sytsma 2011; 2,4) Walker and Sytsma 2007; 3) Drew and Sytsma 2012. | JF301339.1 | DQ667254 | JQ669372 | DQ667443 | JF289068 |
| *Salvia pterocalyx* Hedge. | 3) Will and Claßen-Bockhoff 2017. | / | / | KU578200 | / | / |
| *Salvia pubescens* Benth. | 2,4) Walker and Sytsma 2007. | / | DQ667296 | / | DQ667482 | / |
| *Salvia purpurea* Sessé & Moc. | 2,4) Benitez-Vieyra et al., 2014. | / | MF664636 | / | KJ473981 | / |
| *Salvia pygmaea* Matsum. | 1, 5) Takano and Akiyama 2017; 2) Takano and Okada 2011; 4) Sudarmono and Okada 2007. | LC060854.1 | AB541126 | / | AB295083 | LC060558 |
| *Salvia qimenensis* S.W.Su & J.Q.He. | specimen_voucher="SQIM" | / | JQ934155 | / | / | / |
| *Salvia quezelii* Hedge and Afzal-Rafi | Will and Claßen-Bockhoff 2017. | / | KU563808 | KU578249 | / | / |
| *Salvia radula* Benth. | 1,2) Will and Claßen-Bockhoff 2014. | KJ584293 | KJ584180 | / | / | / |
| *Salvia ranzaniana* Makino. | 2,4) Sudarmono and Okada (2007); 5) Takano 2017. | / | AB287375 | / | AB287374 | LC060560 |
| *Salvia repens* Burch. ex Benth. | Will and Claßen-Bockhoff 2014. | KJ584295 | KJ584231 | KJ747282 | / | / |
| *Salvia retinervia* Briq. | 2,4) Fragoso-Martinez et al., 2017 | / | MF664058 | / | MF664058 | / |
| ***Salvia reuterana* Boiss.** | Iran, TARI (102849). | MK204850 | MK213222 | / | / |  |
| ***Salvia reuterana*.** | Iran, TARI (102842). | MK204849 | / | / | / | / |
| ***Salvia rhytidea* Benth.** | Iran, TARI, (102850). | MK204848 | MK213223 | / | / | MK240129 |
| ***Salvia rhytidea*.** | Iran, Bamonar University (16297). | MK204847 | / | / | / | MK240128 |
| *Salvia ringens* Sm. | Will and Claßen-Bockhoff 2017. | / | KU563810 | KU578213 | / | / |
| *Salvia roborowskii* Maxim. | 2) Walker and Sytsma 2007. | / | DQ667289 | / | DQ667474 | / |
| *Salvia roemeriana* Scheele. | 1,5) Drew and Sytsma 2011; 2) walker and Sytsma 2007. | JF301340 | DQ667211 | / | / | / |
| *Salvia rosifolia* Sm. | Will and Claßen-Bockhoff 2017 | / | KU578209 | / | / | / |
| *Salvia rosmarinus* (L.) Schield.,<br>Handb. Med.-Pharm | 1,5) Drew and Sytsma 2011; 2) Trusty et al. 2004; 3) Drew and Sytsma 2012 4) Walker et al., 2004. | JF30328 | AY506649 | JQ669364 | AY570465 | JF289058.1 |
| *Salvia rufula* Kunth. | 2,4) Fragoso-Martinez et al., 2017. | / | MF622179 | / | MF664063 | / |
| *Salvia rugosa* | 2,5) Walker & Sytsma 2007 | / | DQ667290 | / | DQ667475 | / |
| *Salvia rusbyi* Britton ex Rusby. | 2,4) Walker and Sytsma 2007. | / | DQ667266 | / | DQ667452 | / |
| ***Salvia russellii* Benth.** | Iran, TARI (86040). | MK204846 | MK213224 | / | / | MK240130 |
| *Salvia rypara* Briq. | 2,4) Walker and Sytsma 2007. | / | DQ667266 | / | DQ667452 | / |
| *Salvia rzedowskii*<br>Ramamoorthy | 2,4) Fragoso-Martinez et al., 2017. | / | MF664650 | / | MF664067 | / |
| *Salvia sagittata* Ruiz & Pav. | 2,4) Walker and Sytsma 2007. | / | DQ667446 | / | DQ667260 | / |
| ***Salvia sahendica* Boiss. & Buhse** | Iran, TARI (73990). | MK204845 | MK213225 | / | / | / |
| ***Salvia santalonifolia* Boiss.** | Iran, TARI (102846). | MK204844 | MK213226 | / | / | / |
| *Salvia scabiosifolia* Lam. | Will and Claßen-Bockhoff 2017 | / | KU563811 | KU578237 | / | / |
| *Salvia scabra* L. | Will and Claßen-Bockhoff (2014) | KJ584297 | KJ584233 | KJ747285 | / | / |
| *Salvia scapiformis* Hance | 2,4) Deng et al., 2015 |  | KM886675<br>1 |  | KM886852 |  |
| *Salvia schimperi* Benth. | Will and Claßen-Bockhoff 2014 | KJ584298 | KJ584174 | / | / | / |
| *Salvia schlechteri* Briq, | Will and Claßen-Bockhoff 2014 | KJ584299 | KJ584235 | KJ747286 | / | / |
| *Salvia sclarea* L. | 2) Walker and Sytsma 2007; 3,5) Drew and Sytsma 2012 | / | DQ667222 | JQ669373 | / | JQ669265 |
| ***Salvia sclarea*.** | Iran, TARI (69493). | MK204843 | MK213227 | / | / | MK240131 |
| ***Salvia sclareopsis* Bornm. ex Hedge** | Iran, TARI (3533). | MK204842 | MK213228 | / | / | MK240131 |
| *Salvia scrophularifolia* (Bunge)<br>B. T. Drew | 2,4) Walker and Sytsma 2007; 3) Drew and Sytsma 2012; 5) Drew and Sytsma 2011 | / | DQ667330 | JQ669352 | DQ667519 | JF289051 |
| *Salvia scutellaroides* Kunth. | 2,4) Walker and Sytsma 2007 | / | DQ667327 |  | DQ667516 |  |
| *Salvia sericeo-tomentosa* Rech. F. | Will and Claßen-Bockhoff 2017 | / | KU563822 | KU578238 | / | / |
| *Salvia sessei* Benth. | 2,4) Jenks et al., 2010 | / | HQ418889 | / | HQ418991 | / |
| *Salvia sessilifolia* Baker. | 3) Walker and Sytsma 2007; | KJ584303 | DQ667282 | DQ667467 | / | / |
| *Salvia sinica* Migo. | SS201301 | / | KJ397257 | / | / | / |
| *Salvia somalensis* Vatke. | Will and Claßen-Bockhoff 2014 | KJ584304 | KJ584240 | KJ747311 | / | / |
| *Salvia sonomensis* E. Greene. | 1,4) Walker and Sytsma 2007; 2,5) Walker et al., 2015 | KP853008 | DQ667218 | / | DQ667426 | KP853081 |
| *Salvia sophrona* Briq. | 2,4) Fragoso-Martinez et al., 2017 | / | MF664655 | / | MF664073 | / |
| ***Salvia spinosa* L.** | Iran, TARI (102844) | MK204841 | MK213229 | / | / | MK240132 |
| ***Salvia spiraeifolia* Boiss. & Hohen.** | Iran, TARI (1506) | MK204872 | MK21204 |  |  | / |
| *Salvia splendens* Sellow ex Schult. | 2,4) Fragoso-Martinez et al., 2017 | / | MF622186 | / | MF664076 | / |
| *Salvia stachydifolia* Benth. | 2,4) Walker and Sytsma 2007 | / | DQ667267 | / | DQ667453 | / |
| ***Salvia staminea* Montbret & Aucher ex Benth.** | Iran, TARI (51624) | MK204840 | MK213230 | / | / | / |
| *Salvia stenophylla* Burch. ex Benth. | Will and Claßen-Bockhoff 2014 | KJ584305 | KJ584237 | KJ747260 | / | / |
| *Salvia styphelus* Epling | 4) Jenks et al., 2010 | / | HQ418891 | / | HQ418993 | / |
| ***Salvia suffruticosa* Montbret & Aucher ex Benth.** | Iran, TARI (86522) |  | MK213223 |  |  |  |
| *Salvia summa* A.Nelson. | 1) Will and Claßen-Bockhoff 2014; 2,3) Will and Claßen-Bockhoff 2017. | KJ584307 | KU563874 | KU578164 | / | / |
| *Salvia superba* | voucher="PS0134MT01 | / | FJ546849 | / | / |  |
| ***Salvia syriaca* L.** | Iran, TARI (16699). | MK204839 | MK213232 | / | / | / |
| *Salvia x sylvestris* L. | 2) Will and Claßen-Bockhoff 2017; 3,4) Wang et al., 2013. | / | KJ584177 | KJ747292 | KC414323 | / |
| *Salvia taraxacifolia* Hook.f. | 1,2,3) Will and Claßen-Bockhoff 2014; Walker et al., 2004. | KJ584308 | KJ584228 | KJ747270 | AY570497 | / |
| ***Salvia tebesana* Bunge.** | Iran, TARI (84759). | MK204838 | MK213233 | / | / | / |
| *Salvia texana* (Scheele) Torr. | Will and Claßen-Bockhoff 2017. | / | KJ584199 | KJ747267 | DQ667510 | / |
| *Salvia thermarum* Van Jaarsv. | P. Wester and R. Claßen-Bockhoff 336 (MJG 041398). | KJ584309 | / | KJ747288 | / | / |
| *Salvia thymoides* Benth | 2,4) Walker and Sytsma 2007. | / | DQ667273 | / | DQ667458 | / |
| *Salvia thyrsoflora* Benth. | 2,4) Fragoso-Martinez et al., 2017. | / | MF664668 | / | MF664088 | / |
| *Salvia tiliifolia* Vahl. | 2,4) Fragoso-Martinez et al., 2017. | / | MF664669 | / | MF664089 | / |
| *Salvia tobeyi* Hedge. | Will and Claßen-Bockhoff 2017. | / | KU563865 | KU578188 | / | / |
| *Salvia tomentosa* Mill. | Will and Claßen-Bockhoff 2017. | / | KU563816 | KU578214 | / | / |
| *Salvia tonaticensis*<br>Ramamoorthy ex Lara,<br>Bedolla et Zamudio. | 2,4) Fragoso-Martinez et al., 2017. | / | MF664670 | / | MF664090 | / |
| *Salvia tortuosa* Urb. | 2,4) Jenks et al., 2010. | / | HQ418893 | / | HQ418995 | / |
| *Salvia trichocalycina* Benth. | Walker & Sytsma 2007 | / | DQ667283 | / | DQ667468 | / |
| *Salvia trichoclada* Benth. | Will and Claßen-Bockhoff 2017. | / | / | KU578243 | / | / |
| *Salvia trichostephna* Epling. | 2,4) Fragoso-Martinez et al., 2017. | / | MF664671 | / | MF664091 | / |
| *Salvia tricuspis* Franch. | China: Sichuan | / | EF373633 | / | EU220730 | / |
| *Salvia trijuga* Diels. | 2) Will and Claßen-Bockhoff 2017; 4)<br>Wang et al 2013 | / | KU563779 | / | KC414322.1 | / |
| *Salvia tubifera* Cav. | 2,4) Fragoso-Martinez et al., 2017. | / | HQ418894 | / | HQ418996 | / |
| *Salvia tubiflora* Sm. | 2,4) Fragoso-Martinez et al., 2017. | / | MF664672 | / | MF664092 | / |
| *Salvia uliginosa* Benth. | 2,4) Jenks et al., 2010. | / | HQ418895 | / | HQ418997 | / |
| ***Salvia urmiensis*** Bunge. | Iran, TARI (19586). | MK204837 | MK213234 | / | / | / |
| *Salvia vaseyi* (Porter) Parish. | 4)Walker et al., 2015. | KP853013 | / | / | KP852932 | KP853083 |
| *Salvia veneris* Hedge | Will and Claßen-Bockhoff 2017. | / | KJ584170 | KJ747306 | / | / |
| *Salvia venulosa* Epling | 2,4) Jenks et al., 2010. | / | HQ418896 | / | HQ418998 | / |
| ***Salvia verbascifolia*** M.Bieb. | Iran, TARI (88803). | MK204831 | MK213235 |  | / |  |
| *Salvia verbenaca* L. | Will and Claßen-Bockhoff 2014. | / | KJ584183 | KJ747298 | / | / |
| *Salvia vermifolia* Bedge and<br>Huber-Morath. | Will and Claßen-Bockhoff 2017. | / | Ku563866 | KU578192 | / | / |
| ***Salvia verticillata*** L. | Iran, TARI (2765). | MK204836 | MK213236 | / | / | / |
| ***Salvia virgata*** Jacq. | Iran, TARI (3772). | MK204835 | MK213237 | / | / | / |
| ***Salvia viridis*** L. | Iran, TARI (13283). | MK204834 | MK213238 | / | / | MK240135 |
| *Salvia viscosa* Jacq. | Will and Claßen-Bockhoff 2017. | / | KU563838 | KU578186 | / | / |
| *Salvia vvedenskii* Nikitina . | 2,3) Will and Claßen-Bockhoff 2017. | / | KU563879 | KU578202 | / | / |
| *Salvia whitehousei* Alziar | 1,2,3) Will and Claßen-Bockhoff 2014; 4) Walker and Sytsma 2007. | KJ584311 | KJ584198 | KJ747268 | DQ667509 | / |
| ***Salvia xanthocheila*** Boiss. ex Benth. | Iran, TARI (69728). | MK204833 | MK213239 | / | / | MK240136 |
| *Salvia yangii* B. T. Drew | 1,5) Drew and Sytsma 2011; 2) Will and Claßen-Bockhoff (2014); 3) Drew and Sytsma 2011. | JF301328 | KJ584242 | JQ669352 |  | JF289051` |
| *Salvia yunnanensis* C.H.Wright. | 2) YunN0309-2; 3) G.X. Hu & al., QT001, 4) H.F. Guo 2017257(PE) | / | DQ132866 | KT210283 | EF014356 |  |

### 2.2. DNA extraction, amplification, and sequencing

Total DNA was extracted from herbarium and silica-dried material using a modified CTAB method (Doyle & Doyle 1987) in which, to break down secondary metabolites, the mixture of ground leaf tissue and CTAB solution was kept at room temperature for 24 hours. ITS, ETS, and *ycf1* regions were amplified using the polymerase chain reaction (PCR) with each sample prepared in 25-μl volumes with the following components: 1 μl of DNA solution (20 ng), 2.5 μl of reaction buffer, 2 μl dNTP mix (0.2 mM), 1 μl of each primer (10 uM), 1 μl of MgCl2, and 1.5 μl of *Taq* DNA polymerase. The PCR conditions for the nuclear regions for most species were: 95°C for 2 minutes, 32 cycles of denaturation for 20 seconds at 94°C, primer annealing for 20 seconds at 50°C, and 2 minutes extension at 72°C, with a final extension of 7 minutes at 72°C. For the *ycf1* region, we modified the annealing temperature to 52°C for 1 minute (PCR optimization was set based on personal communication with B. Drew). High-quality PCR products were sequenced on an ABI 3730 DNA Analyzer (Applied Biosystems, Inc.) at the University of Florida Interdisciplinary Center for Biotechnology Research (ICBR).

### 2.3. Alignment and phylogenetic analysis

All consensus DNA sequences were generated using Geneious Pro v. 10.22 (Biomatters, Auckland, New Zealand). Alignments were performed with the MAFFT plugin in Geneious with manual adjustment. Maximum likelihood analysis was performed using the CIPRES Science Gateway with RAxML HPC v.8 on XSEDE using the GTRGAMMA model with _Fa (rapid bootstrapping analysis/search for the best ML tree) with 1000 iterations for bootstrapping. Default settings were used for other options. Phylogenetic analyses were conducted for 1) all plastid loci (*rpl32-trnl*, *trnl-trnf*, and *ycf1-rps15*), 2) both nuclear loci (ITS and ETS), and 3) a combined data set of plastid and nuclear loci.

### 2.4. BEAST analysis (divergence time estimation)

We estimated divergence times using BEAST version 2.2.0 (Bouckaert et al., 2014) under the uncorrelated lognormal model. Priors for the branch rate were assumed as a Yule process. A node prior was calibrated for the most recent common ancestor (MRCA) of *Melissa* and *Lepechinia* (28.4 Ma with a mean of 1.5 and a SD of 0.5; Drew and Sytsma 2012; Kriebel et al., 2019) with a lognormal distribution. The BEAST analysis was performed with two independent runs of Markov Chain Monte Carlo. Each run was performed for 2*10^8^ generations, with parameters logged every 1000 generations. We used Tracer v. 1.6 to evaluate the ESS (Effective Sample Size) to assure that the chains were run sufficiently long. An ESS > 200 indicates that the two independent runs were adequate. Tree Annotator was used to find the maximum clade credibility reporting median node ages after discarding the first 10% of the generations as burn-in.

### 2.5. Ancestral state reconstruction

Two characters with discrete states were scored: mode of lever mechanism (present / absent) and habit (woody / herbaceous). We treated shrubs and subshrubs as woody; however, distinguishing woody from herbaceous is not always straightforward because some mostly herbaceous plants may become woody in special climatic situations (FitzJohn et al., 2014; Zanne et al., 2014). Therefore, we treated a species as woody if it is considered a shrub or subshrub in the literature or if it was defined as having a woody rootstock. In addition, the continuous character corolla length was measured from the joint of the calyx to the end of the upper lip.

The relevant data for the discrete and continuous traits were collected from the literature: *Flora of China* (www.efloras.org/flora_page.asp? flora_id = 2), *California Salvia* (Epling, 1983), *Flora of USSR* (Pobedimova, 1954), *Flora of Turkey and the East Aegeans* (Hedge, 1982), *Flora Iranica* (Hedge, 1982), *Flora of Southern Africa* (Codd, 1985), *Flora of Madagascar* (Hedge, 1992), *Flora dels Paiso Catalans* (Bolos and Vigo, 1995; Wester and Claßen-Bockhoff, 2011), and *Flora of Iran* (Jamzad, 2012). Additionally, we used online resources (www.gbif.org; www.tropicos.org) as sources of data. For some species, the corolla length was measured using the digitized type specimen available on JSTOR’s Global Plants database (<u>http:/plants.jstor.org</u>).

For the discrete data, we used maximum likelihood to define the best model fitting our data using the function ‘ace’ implemented in the R package ape v5. 3 (Paradis et al., 2004). We tested “ER” (Equal Rates) and “ARD” (All Rates Different) on our data, and the best model was selected based on the Akaike Information Criterion (AIC) (Akaike, 1974). We used the Akaike weight using aic.w function in the R package geiger v2.0.6 to select the best model for those data wih low delta AIC between ER and ARD models. To reconstruct ancestral states, we used stochastic character mapping with 1000 iterations using the make.simmap function in the phytools v0.7.78 package (Revell, 2012). We also reconstructed the ancestral state of corolla length using the lik.anc in phytools to calculate the likelihood of each ancestral state. Ancestral states of corolla length and 95% confidence intervals were evaluated using the function anc.ML with an OU (Ornstein-Unlenbeck) model in phytools.

### 2.6. Macroevolutionary patterns within corolla length

We focused on corolla length as one of the most important morphological traits that might influence pollinator-flower interactions (Fernández- Mazuecos et al., 2013; Gómez et al., 2016; Landis et al., 2018). To investigate the evolutionary dynamics of corolla length throughout *Salvia* phylogeny, we applied three quantitative approaches based on the time-calibrated phylogeny as follows:

#### 2.6.1. Diversification model

We examined three evolutionary models with different patterns of phenotypic evolution using the R package geiger v2.0.6 (Harmon et al., 2008) following three different models. 1) The Brownian Motion model (BM): This model describes a “random walk” of evolution for continuous characters. 2) The Ornstein-Uhlenbeck model (OU): This model describes the local occurrence of stabilizing selection in which the trait is drawn toward optimal fitness (Hansen 1997). 3) The Early Burst model (EB): This model is known as a classic model of adaptive radiation, in which the initial stage of morphological evolution is rapid with decreasing morphological evolution after ecological spaces are filled (Harmon et al., 2010). Based on a recent model of diversification reconstructed by Aguilée et al. (2018), after an initial phase of geographic adaptive radiation, diversification rates can be affected not only by ecological niches, but also by genetic processes, competition, and landscape dynamics. The best model for explaining diversification of *Salvia* was selected based on the AIC (Akaike, 1974).

#### 2.6.2. Disparity Through Time

Disparity Through Time (DTT) of the corolla length was modeled using the R package geiger v2.0.6 (Harmon et al., 2008). This analysis uses corolla length of extant *Salvia* species to reconstruct ancestral corolla length values and model disparity between species. This approach estimates the pairwise Euclidean distance of the trait over time and compares it with the expected value under a null model of Brownian motion by iterative simulation.

Phenotypic disparity refers to the phenotypic variation among related species (Harmon, 2003). We simulated corolla length evolution with 10,000 generations across the phylogenetic tree built from the combined data set of plastid and nuclear sequences. The Morphological Disparity Index (MDI) was calculated, and the average disparity of corolla length from the real and simulated data was plotted. Negative MDI shows lower disparity of the trait than expected, and positive MDI indicates strong overlap in morphospace and higher disparity within subclades (Donoso et al., 2015).

#### 2.6.3. iversification rate

To assess variation in rates of diversification of corolla length across *Salvia*, we used the phenotypic trait module in BAMM. We simulated 20,000,000 generations, and the priors were set using the function “SetBAMMpriors” in the R package BAMMTools v.2.1.6 (Rabosky et al., 2014). We specified the sampling fraction by accounting for the number of samples for each of the four major clades. Sampling fractions were set as: 0.47 (clade I), 0.26 (clade II), 0.95 (clade III) and 0.56 (clade IV). We performed MCMC simulation with 20,000,000 generations by sampling every 1000 generations. We discarded the first 25% of runs as burn-in. Effective Sample Size (ESS) > 200 was used to evaluate the convergence of four Markov Chain Monte Carlo chains. The BAMM output was analyzed using BAMMtools.

### 2.8. Lever-mechanism-dependent diversification

To examine whether the diversification rate in *Salvia* is correlated with the presence of the lever mechanism, we applied HiSSE (Hidden State Speciation and Extinction) implemented in the R package hisse v2.1.1 (Beaulieu and O^’^Meara 2016), which is a modified method of BiSSE (Binary State Speciation and extinction) (Maddison et al., 2014). Rabosky (2014) argued that the BiSSE method suffers from type Ι and type II errors. In those cases, traits that are not biologically correlated with speciation rates show significant effects on diversification (Goldberg and Rabosky, 2015). In other words, rejecting the null hypothesis in BiSSE does not mean the alternative is true.

Compared with the BiSSE model, the HiSSE model considers more free parameters and assumes a hidden state for each of the observed states that potentially have independent rates of diversification (0A, 1A, 0B, 1B). The Character Independent Diversification (CID) models, which assume independent evolution for binary characters, were also implemented. The CID models explicitly test that the evolution of a binary character is independent of the diversification process without forcing the diversification process to be constant. Different subsets of the HiSSE model that differ in speciation, extinction, and transition rate parameters, along with standard BiSSE models, were estimated (cf. Harrington and Reeder, 2017). We accounted for incomplete taxon sampling in our phylogeny by assigning the sampling frequency of each state as 0.256 (presence of the lever mechanim) and 0.056 (absence of the lever mechanism). The model average of ancestral state and diversification of all fitted models was plotted using the function “plot.hisse.state”. The advantage of this function is that it accounts for both state and rate uncertainty of the models along plotted branches. We also used FiSSE (Fast intuitive State-dependent Speciation Extinction) as a non-parametric test for the lever-mechanism-dependent speciation rate. This method does not depend on the character state, but considers the distribution of branch lengths (Rabosky and Goldberg 2017).

### 2.9. Diversity-dependent diversification

We also used the R package DDD v2.7 (Etienne and Haegman, 2012) to test whether diversification in *Salvia* is dependent or independent of diversity. DDD uses a hidden Markov model to calculate the likelihood of phylogenetic history under a diversity-dependent birth-death model of diversification. DDD estimates “*K*”, the maximum number of species that a clade can have in a given environment; a value of *K* near the number of extant species suggests that a clade is close to its ecological limit. The other two models are a density-dependent logistic (DDL+E) model and a density-dependent exponential (DDE+E) model. The model with the lowest AICc was selected as the best model. We also calculated the maximum likelihood evolutionary history pattern with both a Yule model and a constant rate birth-death (CrBD) model. The Maximum Clade Credibility (MCC) of the BEAST output was used to perform this analysis. We also examined four model fits as an alternative method using the R package laser v2. 4 (Rabosky 2006).

## 3. Results

### 3.1. Characteristics of the phylogenetic data matrix and phylogenetic analysis

In this study, 143 new DNA sequences were generated for 50 Iranian *Salvia* species, including 50 species (59 accessions) for the ETS region, 46 species (47 accessions) for ITS, and 34 species for *ycf1*.

Maximum likelihood analyses of all three data combinations were conducted: 1) plastid loci (*rpl32-trnl*, *trnl-trnf*, and *ycf1-rps15*), 2) nuclear loci (ITS and ETS), and 3) a combined data set of plastid and nuclear loci. The overall topologies of the nuclear loci, plastid loci, and the combined data set are smilar in recovering major clades. Based on both nuclear and plastid regions, the phylogenetic relationships among most of the Eurasian species in clade I are unresolved. This result is not surprising given that we had more missing data in the plastid partition than other partitions. The nuclear and combined data sets provide higher support for most of the clades than plastid data. For example, clade III was recovered as fully resolved based on nuclear regions and combined data, but based on plastid data, the relationship between *S. majdaea* and *S. macilenta* with the remain group in clade III was unresolved. Trees based on the nuclear and combined data sets were highly similar, with only some minor differences in support values for terminal clades. However, the combined data recovered a more resolved phylogney. For example, in clade I within Subgen. *Heterosphace*, the relationship between the *S. verticillata* group with the remaining taxa was resolved based on the combined data but not the nuclear data. As a result, we used the results from the combined data set in subsequent analyses and in our discussion below.

We recovered four major clades of *Salvia* species: clade I (Eurasian and southern African *Salvia)*, clade II (South and North American *Salvia*), clade III (southwestern Asian and northern African *Salvia*), and clade IV (Southeast Asian *Salvia*). With more taxon sampling, we provide a new phylogeny for clade III with species that are primarily distributed in southwestern Asia. However, we mostly focus here on newly recovered relationships for Iranian species of *Salvia* and the clades to which they belong rather than on *Salvia* phylogeny as a whole. For more straightforward comparisons, we also used both clade names provided in previous phylogenetic studies of *Salvia* (Will and Claßen-Bockhoff, 2017) along with the recent classification (Drew et al., 2017; Kriebel et al., 2019) in **Suppl. 1.**

#### Clade I

In this clade, species are mainly distributed in Europe, Central Asia, western Asia, and southern Africa. This clade contains 140 species (170 accessions) out of the 250-300 species described for these areas. They fall into four distinct subclades (subclades I-A, I-B, I-C, and I-D) of Will and Claßen-Bockhoff (2017) including subgenera *Salvia* Benth., *Sclarea* Benth., and *Heterosphace* Benth (Kriebel et al., 2019). For the first time, resolution within the *S. verticillata* group was obtained. *Salvia taraxacifolia* was recovered as the sister to the *S. verticillata* group (BS = 81%) consisting of *S. verticillata*, *S. judaica*, and *S. russellii.* This clade was in turn sister (BS = 95%) to the southern African clade.

Subclade I-C and I-D: 26 species of Iranian *Salvia* sampled here were recovered as members of subclades I-C, and six species were placed in subclade I-D, including subgen. *Sclarea* and *Salvia* based on the updated subgeneric classification of *Salvia* (Will and Claßen-Bockhoff 2017; Drew et al., 2017; Kriebel et al., 2019). These subclades include *Salvia* distributed in western Asia (Afghanistan, Iran, Iraq, and Turkey), Central Asia, Europe, and the Canary Islands. Based on the combined data set of nuclear and plastid loci, the phylogenetic relationships among most of the taxa were unresolved. However, several groups were identified within this polytomy: 1) *S. jamzadaei, S. macrochlamys*, *S. bracteata*, and allied species; 2) two species endemic to Iran (*S. leriifolia* and *S. hypochionaea*) along with *S. montbretia*, *S. daghestanica*, and *S. phlomoides*; 3) *S. spinosa*, *S. sclareopsis*, *S. macrosiphon*, *S. reuterana*, *S. perspolitiana*, and *S. palaestina* (a clade with BS = 60%); and 4) *S. nemorosa* and *S. virgata* from southwestern Asia, the Caucasus, and Europe along with *S. deserta* and *Salvia × sylvestris*.

#### Clade II

*Salvia* species in this clade are endemic to South and North America, and clade II includes more than half of all *Salvia* species, with approximately 600 species in subg. *Calosphace* and 19 species in subg. *Audibertia*. Clade II is recovered with BS= 97%; however, relationships among taxa in this clade are not fully resolved. Clade I is sister to clade II with BS= 74%.

#### Clade III

This clade contains species of *Salvia* from northern Africa and southwestern Asia. We provide the most comprehensive taxon sampling for clade III to date by generating 13 new sequences for members of this group from the region of Iran. Clade III was recovered with 100% BS support with a well-resolved phylogeny in trees from nuclear and combined data. Based on the combined nuclear and plastid tree, *S. majdae*, which was placed in subgen. *Zhumeria* (Drew et al., 2017), was found instead to be sister with high support (BS = 97%) to the *S. aristata* group, which includes *S. pterocalyx* (from northeastern Afghanistan) and *S. vvedenskii* and *S. margaritae* (from Central Asia). *Salvia majdae* and the *S. aristata* group were placed as the sister clade to a trichomy of *S. aegyptiaca*, *S. macilenta*, and *S. eremophila*.

#### Clade IV

*Salvia* species in this clade are restricted to eastern Asia, with the exception of *S. glutinosa* and *S. plebeia.* We provide new sequence data for *S. glutinosa*, which is distributed in the northern part of Iran and some parts of Europe. *Salvia plebeia* is reported from Iran and Afghanistan and extends to Southeast Asia. Based on our results, *S. glutinosa* forms a clade with *S. nubicola*, *S. koyamae*, *S. glabrescense*, and *S. nipponica* with BS = 88%.

### 3.2. Divergence times

Divergence times were estimated using the combined data set of nuclear and plastid loci. The results are congruent with previous results (Drew et al., 2017; Kriebel et al., 2019). Our BEAST analysis (**Fig. 3**) suggests that *Salvia* originated in the Oligocene ∼34 Ma. Divergence time estimation showed that the split between clade I and the rest of *Salvia* occurred approximately 31 Ma (95% HPD = 37.6-27.5 Ma). In clade I, the North American clade diverged from the African and Mediterranean clade approximately 15 Ma (95% HPD = 20.06-10.69) during the middle Miocene. The age of the MRCA of most of the Iranian *Salvia* species in clade I was estimated as 14.3 Ma (95%HPD = 18.74-10.47 Ma) near the end of the early Miocene. Clade II diverged from clade III (mostly from southwestern Asia) in the early Miocene (95% HPD = 27.8-17.8 Ma). The split between clade IV (eastern Asia) and clade III (southwestern Asia and northern Africa) is estimated to have occurred during the late Oligocene (95%HPD = 31.5-20.26 Ma).

**Fig. 3:**
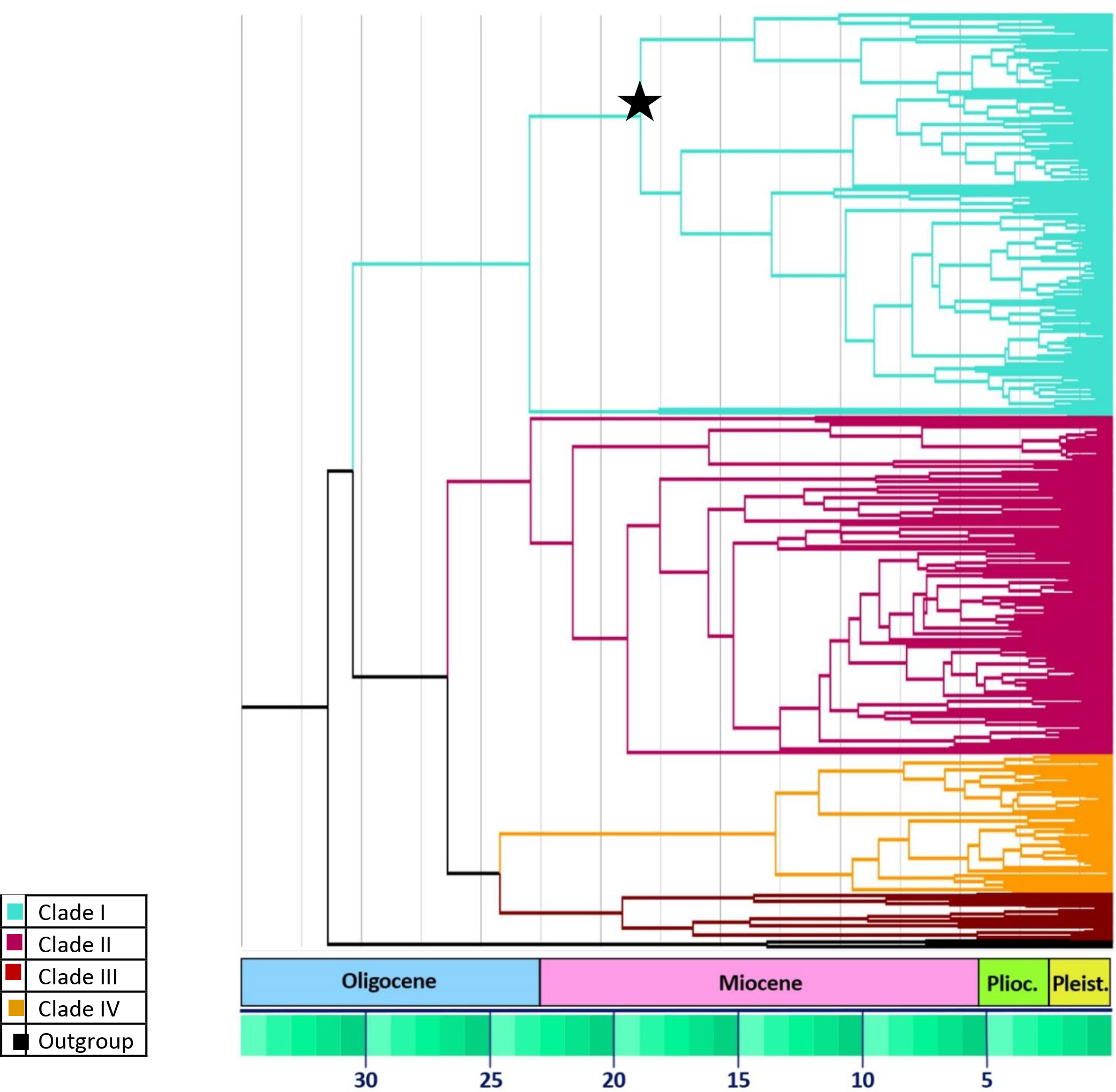
Maximum Clade Credibility (MCC) obtained from BEAST analysis based on five combined nuclear and plastid spacer regions. The map is colored based on four identified clades in *Salvia*. The *x*-axis represents the age range of extant *Salvia* lineages. The star indicates divergence of the southwestern Asia clade I (Turkey and Iran) including subgenera *Sclarea* and *Salvia* from Subgenus *Heterosphace*

### 3.3. Ancestral character reconstruction

Corolla length varies from 4 mm in *S. aegyptiaca* in clade III to 51 mm in *S. patens* in clade II. The most recent common ancestor for *Salvia* was reconstructed as having a corolla length of approximately 15-18 mm (**Fig. 4)**. Corolla length of ∼20 mm was inferred as the ancestral state for clade I. In clade I, within subg. *Sclarea*, multiple shifts from a corolla length of 20 mm to smaller corollas occurred, but in subg. *Salvia*, all the species evolved corolla lengths longer than 20 mm. In subg. *Heterosphace*, including bird-pollinated species of *Salvia* from southern Africa (*S*. *africana*-lutea, *S. lanceolata*, *S. thermarum*), the corolla length is more variable, ranging from ∼7-41 mm. In clade II, shifts in the range of corolla size were much higher than in other clades, especially in subg. *Calosphace*. In clade III, the ancestral state of corolla length was recovered as ∼15 mm. Within this clade, species of *Salvia* have small flowers (4-9 mm) with shifts to larger flowers in subg. *Zhumeria* and the *S*. *aristata* group.

**Fig. 4:**
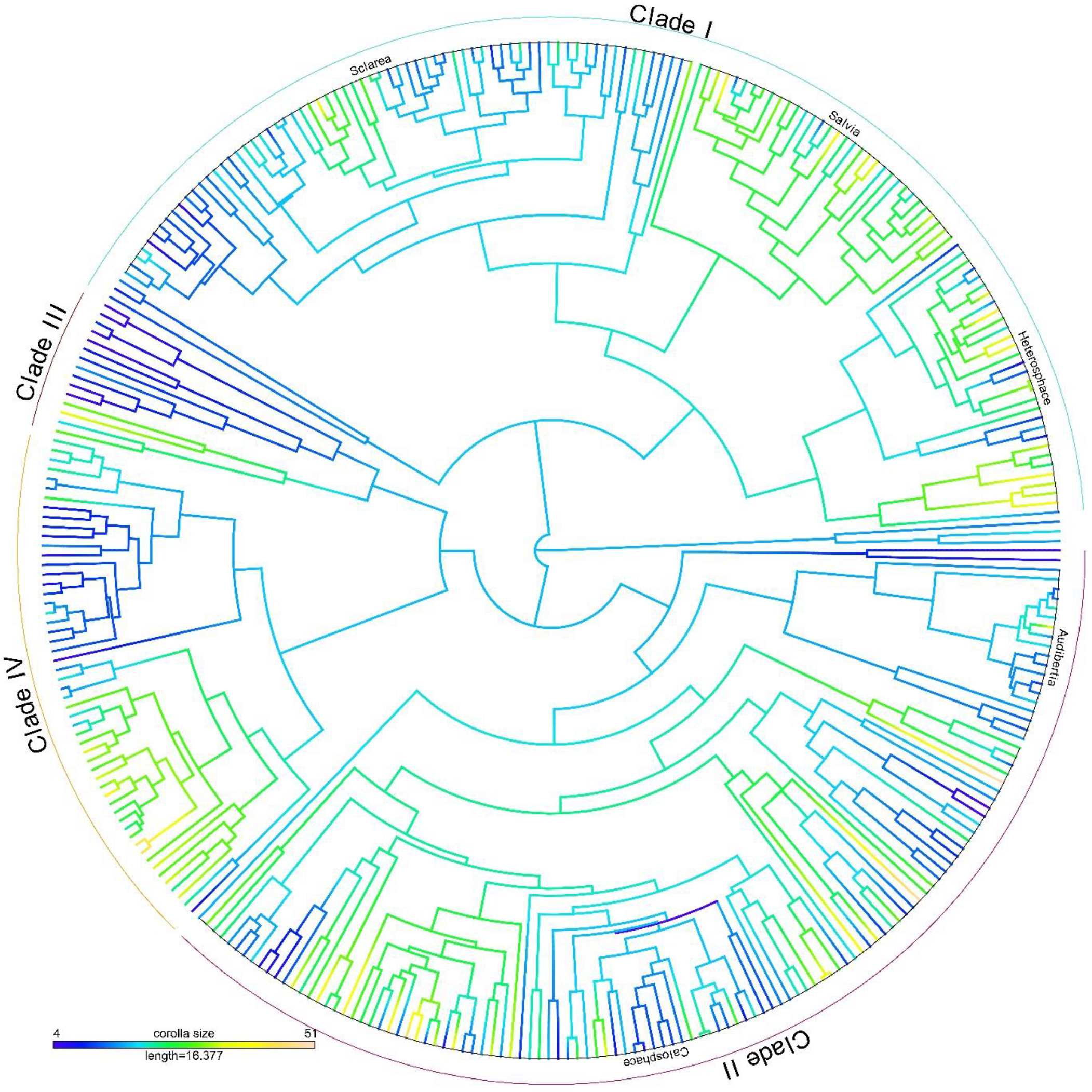
Ancestral reconstruction of corolla length in *Salvia* on a dated phylogeny using maximum likelihood in the phytools R package. The legend indicates the range of corolla length in mm by branch color in *Salvia*. Four distinct clades in *Salvia* are identified by relevant colors on the circumference of the tree.

The ER model was selected as the best model for the evolution of the lever mechanism based on the AIC value. However, the difference between the ER (Equal Rates) and the ARD (All Rates Different) model was minimal (ER 168.46 vs. ARD 169.68). The Akaike weight for the ER model (0.65) was higher than for the ARD model (0.32). Therefore, we reconstructed the ancestral state of the lever mechanism based on the ER model. The ancestral state of the lever mechanism for *Salvia* was equivocal **(Fig. 5**). In clade I, the ancestral state of the lever mechanism was also equivocal, but the ancestral state of subg. *Salvia*, *Sclarea*, and *Heterosphace* is an active lever mechanism. In clade II, the ancestral state is equivocal with several shifts in subg. *Calosphace* from an active lever to a non-active lever. In clade III, *Salvia* species lack an active lever mechanism, and the ancestral state for the clade is equivocal.

**Fig. 5:**
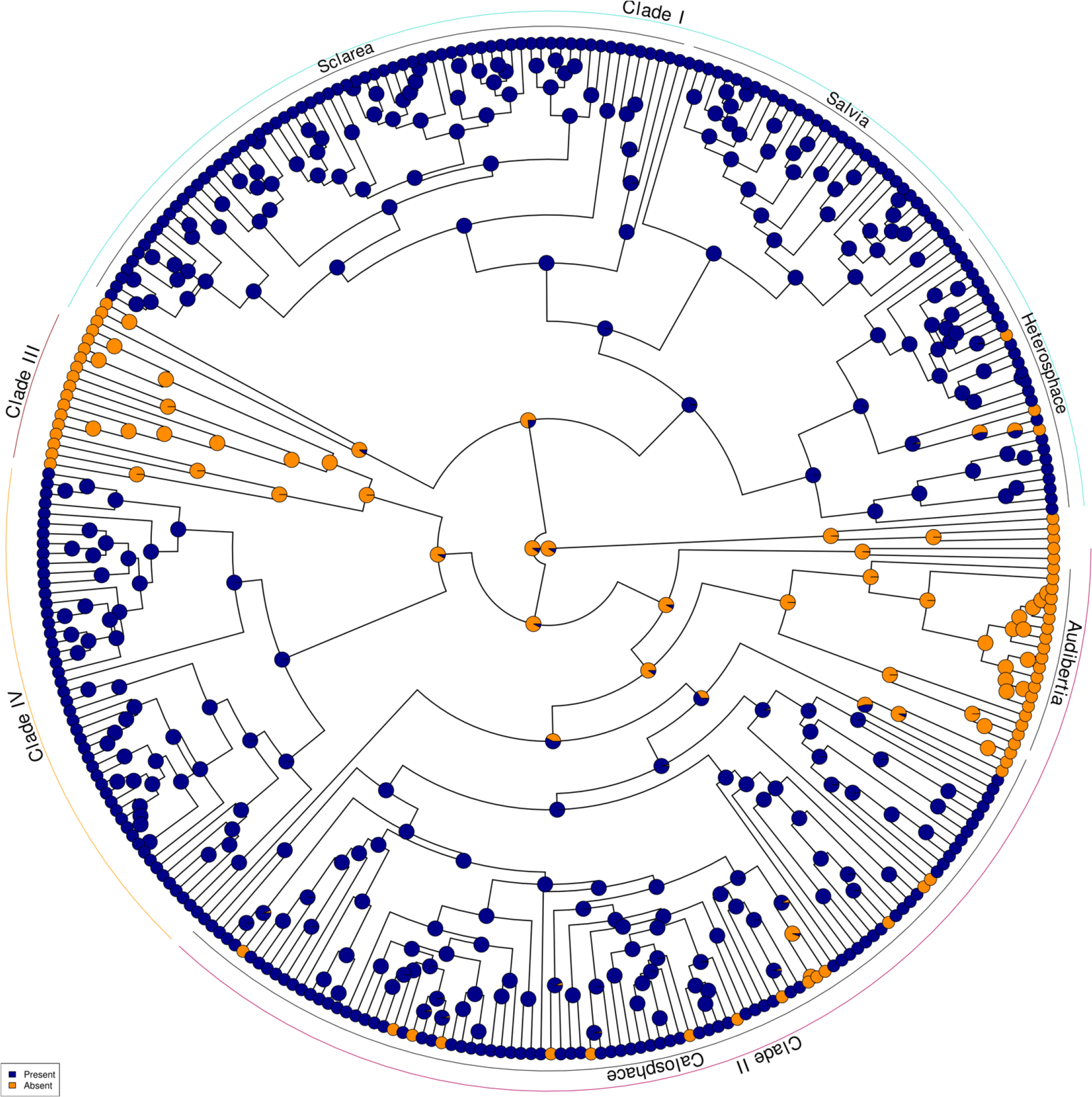
Ancestral reconstruction of the lever mechanism trait (present/ absent) across *Salvia* phylogeny based on likelihood state with ARD model.

The ARD model was moderately suggested as the best model (Delta AIC = 3.345) for inferring the ancestral state of habit across the *Salvia* phylogeny. The ancestral state of habit for all of *Salvia*, as well as major subclades, was found here to be equivocal (**Fig. 6**). In clade I, species of subgen. *Sclarea* are mostly herbs with a few shifts to shrub forms, but subgen. *Salvia* and *Heterosphace* are mostly shrubs with several shifts to herbaceousness. In clade II, especially in subgen. *Calosphace*, shifting from herb to shrub was more frequent than in other clades.

**Fig. 6:**
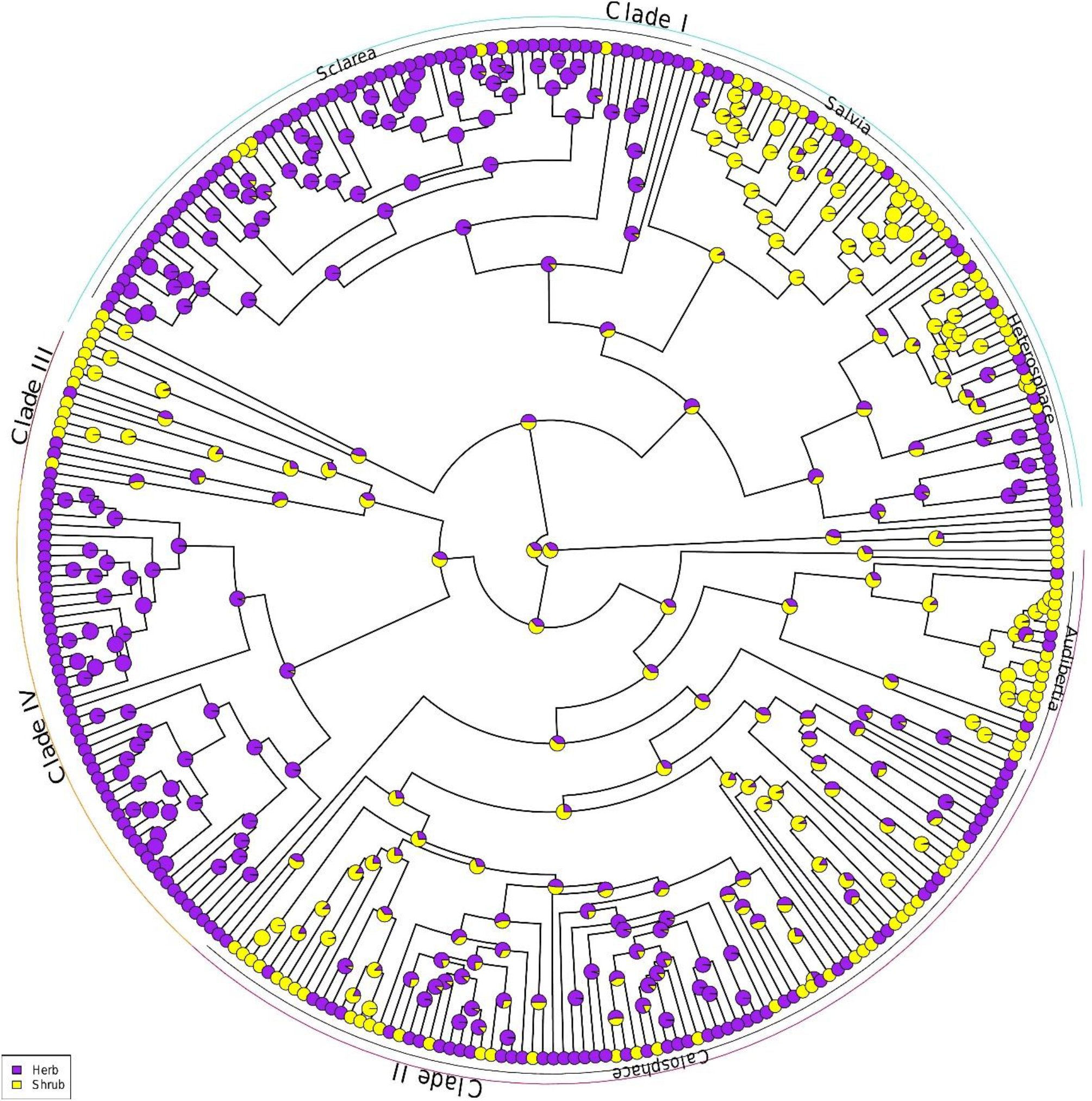
Ancestral reconstruction of habit (herb/shrub) across *Salvia* phylogeny using stochastic mapping in phytools.

### 3.4. Tempo and mode of corolla length evolution

The analysis of disparity through time showed that the rates of diversification in corolla length among subclades of *Salvia* are higher than expected under a null hypothesis of Brownian motion (MDI = +0.21). Therefore, *Salvia* subclades have diversified greatly in corolla length. The corolla length decreased during the Miocene between approximately 17-15 Ma (within the 95% CI calculated from simulations of corolla length disparity), but showed a remarkable increase during the last ∼10 Ma, during which the relative disparity of corolla length is higher than the 95% DTT range of simulated data (**Fig. 7**).

**Fig. 7:**
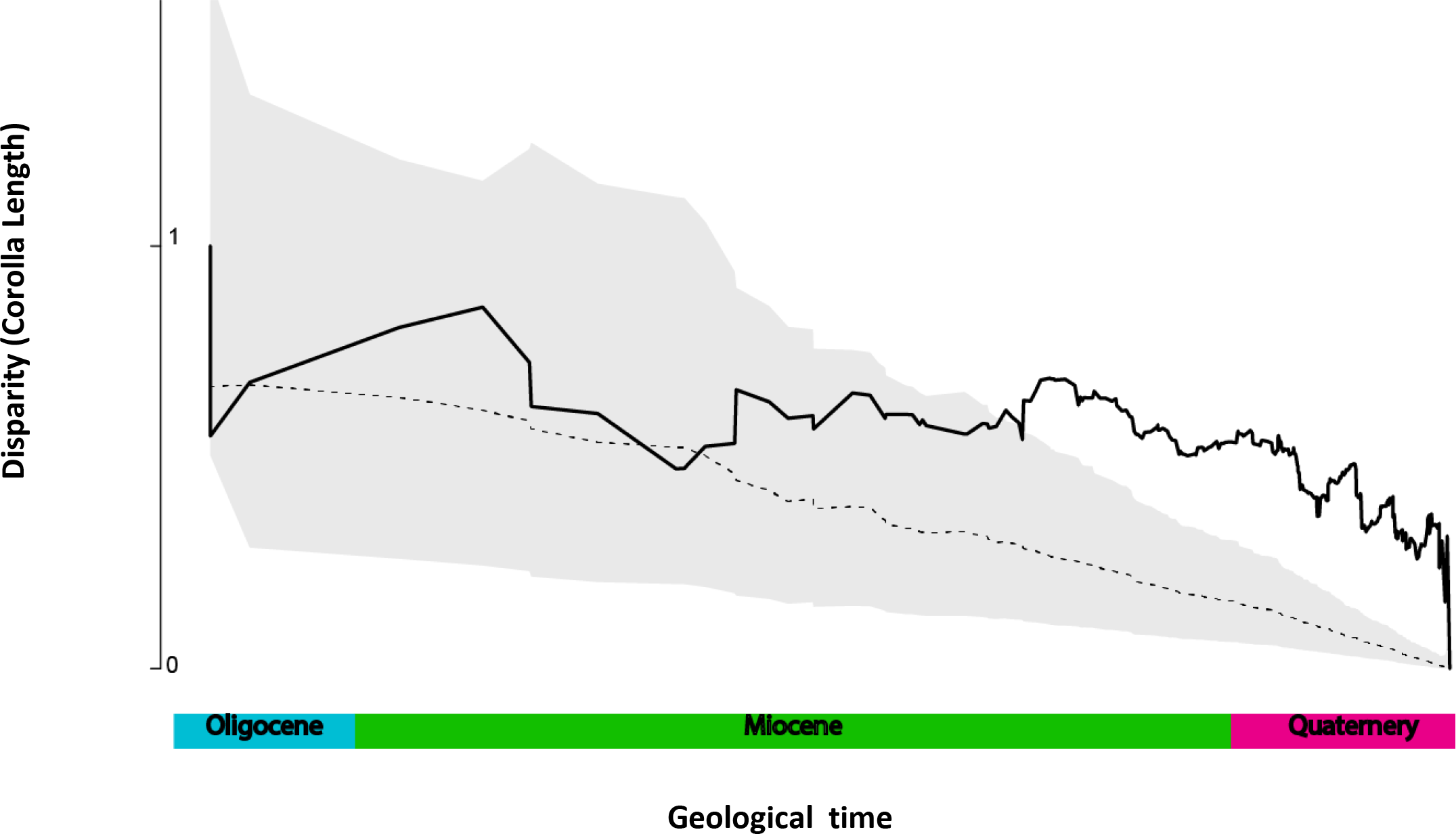
The mean subclade Disparity Through Time (DTT) for corolla length compared with the median subclade disparity under a Brownian motion model. The solid line shows the observed disparity, and the dashed line is the mean disparity of 1000 simulations of corolla length disparity over the phylogenetic tree. The grey shade indicates the 95% confidence interval of DTT.

The model-based analysis of corolla length diversification determined “OU” as the best approximation model of this trait across *Salvia* phylogeny. Hence, our results suggest that corolla length evolution underwent stabilizing selection towards a median value (**Table 2**).

**Table 2:** Rate and modes of Salvia corolla length diversification. Fitted models are Brownian motion (BM), Ornstein-Uhlenbeck (OU) and Early Burst (EB). The best-fit model is estimated based on the lowest bias-corrected Akaike Information Criterion.

| Model | Model Parameters | LogL | AICc | ΔAIC |
| --- | --- | --- | --- | --- |
| BM | $\alpha = 0$ | -1233 | 2470 | 68 |
| OU | $\alpha = 0.15$ | -1198.020364 | 2402 | 0 |
| EB | $\alpha = -0.00001$ | -1233.21 | 2472 | 70 |

#### 3.4.1. Corolla length evolutionary rates

To assess whether the MCMC output of the BAMM analysis for corolla length has converged, we checked the effective sample sizes of the log-likelihood and the number of shift events. Based on ESS_Number of shifts_ = 1139.461 and ESS_Loglike_= 1458.028, the MCMC simulation converged. The phylorate plot confirmed heterogeneous rates of evolution of corolla length in *Salvia*. The best distinct shift configuration with the highest posterior probability was detected at the MRCA of core *Calosphace* in clade II (**Fig. 8**).

**Fig. 8:**
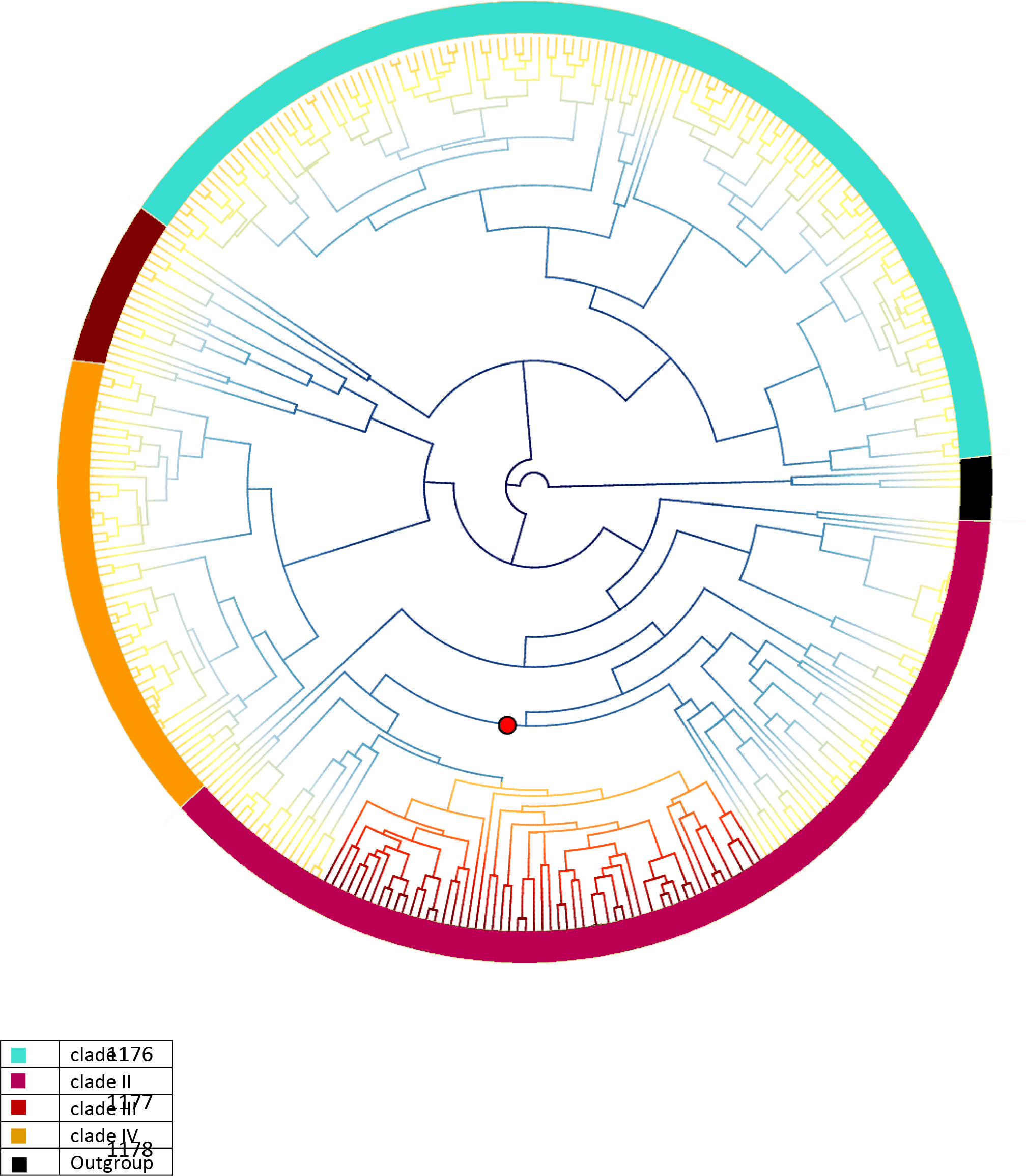
Corolla length evolution across *Salvia* phylogeny based on a BAMM analysis. The best shift was detected in clade II in *Calosphace* clade

### 3.5. Diversity-dependent diversification

The maximum likelihood analysis of lineage diversification showed that among four fitted models, with two models dependent on diversity, the Yule model was selected as the best model based on AIC values for explaining *Salvia* diversification through time. Hence, the evolutionary pattern of S*alvia* diversification is independent of diversity **(Table 3**). The estimated carrying capacity (*K>3000*), which refers to the potential number of species that a clade can sustain, is higher than the number of extant species of *Salvia* (∼1000). Rejection of the diversity-dependent diversification model implies that *Salvia* has not reached its ecological limit in terms of number of species and that speciation has not yet started to decline due to increased species competition or fewer ecological resources.

**Table 3:** Rates of *Salvia* diversification examining multiple evolutionary models. Models fitted include diversity-dependent linear speciation and extinction (DDL + E), diversity-dependent exponential speciation and extinction (DDE+E) and two constant-rate diversification models: a pure-birth (Yule) model and birth-death (crBD) model. λ = speciation rate (Ma/lineage); µ = extinction rate (Ma/lineage); *K* = carrying capacity; AIC = Akaike Information Criterion (AIC) for testing model fit. The capacity for the potential number of *Salvia* species is higher than the number of extant species (∼1000 spp.), suggesting that current *Salvia* diversification is independent of diversity.

| Model | lambda | mu | K | loglik | AIC |
| --- | --- | --- | --- | --- | --- |
| Yule | 0.246 | 0 | ----- | -989.739 | 1977.782 |
| CrBD | 0.290 | 0.079 | ----- | -987.888 | 1979.768 |
| DDE+E | 0.6682 | 0.2331 | 36504.2 | -993.5864 | 1993.169 |
| DDS+E | 0.3579 | 0.14616 | 3315.75 | -988.042 | 1982.86 |

### 3.6. Lever-mechanism-dependent diversification

We used HiSSE (Beaulieu and O^’^Meara, 2016) and FiSSE (Rabosky and Goldbeg., 2017) methods for analyzing the effect of the active lever mechanism on diversification. We found that the HiSSE model with an equal irreversible transition rate among states (q0B1B=0, q1B0B=0, all other q’s equal) is the best model for explaining the effect of the lever mechanism on *Salvia* diversification. Better performance of the HiSSE model than the BiSSE model indicates a signal of lever-mechanism-dependent diversification as well as a signal of other unobserved or unmeasured traits. Therefore, we infer that the lever mechanism is indirectly responsible for *Salvia* diversification (**Table 4**). Based on FiSSE two-tailed parameters, the P-value = 0.69; therefore, the null hypothesis of a close association between the lever mechanism and *Salvia* diversification is rejected. The average tip rate of diversification for an active lever mechanism is λ 1 = 0.26 and for a non-active lever mechanism is λ0 = 0.22.

**Table 4:**
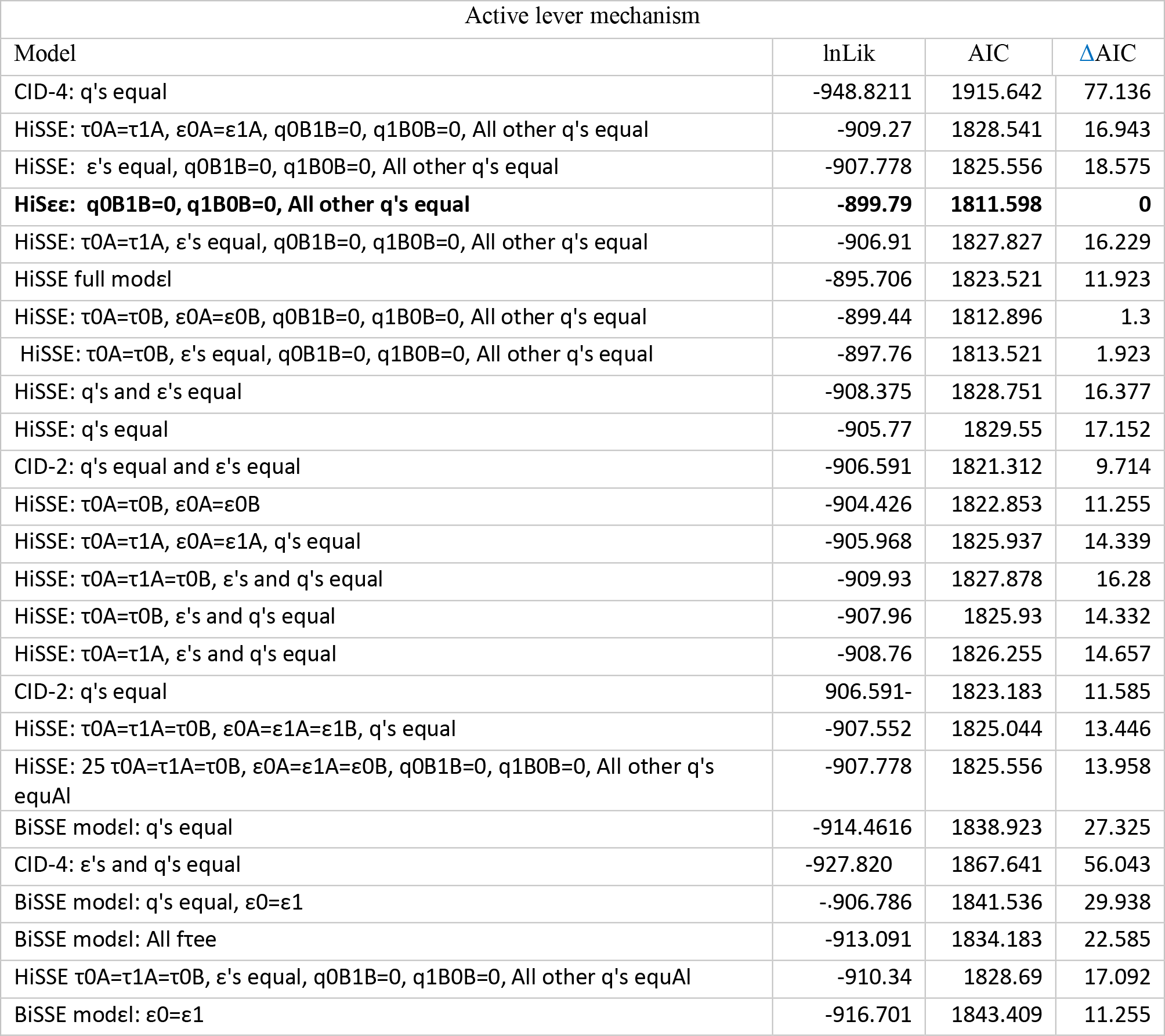
HiSSE model subsets that were fitted for study of the effect of the lever mechanism on diversification across Salvia phylogeny. The best-fit model shown in bold was selected based on a bias-corrected Akaike Information Criterion (AICc).

## 4. Discussion

### 4.1. Phylogeny

By including 50 species of *Salvia* from the Iran region, the limited taxon sampling for the Eastern Hemisphere encountered in previous studies (Drew and Sytsma, 2012; Will and Claßen-Bockhoff, 2014, 2017; Drew et al., 2017) was remedied to some extent. To reconstruct the phylogenetic tree of *Salvia* as comprehensively as possible, our newly generated data were combined with relevant sequences from previous studies (Walker and Sytsma, 2007; Drew and Sytsma, 2012; Takano and Okado 2011; Will and Claßen-Bockhoff, 2014; Fragoso- Martineze, 2017).

Our results for *Salvia* as a whole are similar to those reported in previous phylogenetic results in recovering four major well-supported clades comprising six subgenera were recovered for *Salvia* (Will and Claßen-Bockhoff 2017; Drew et al., 2017; Kriebel et al., 2019). The resultant trees for all the data sets largely agree with each other. Nevertheless, some discrepancies in phylogenetic relationships within subclades were observed. In the paragraphs that follow, we summarize and discuss the major phylogenetic results of this study and also compare our findings to other studies of *Salvia*.

#### Clade I

This clade includes species from three subgenera, *Heterospahce*, *Salvia*, and *Sclarea* (Drew et al., 2017; Kriebel et al., 2019). *Salvia* species in this clade are distributed in small areas of North America, southern Africa and Madagascar, western Asia, Europe, and the Canary Islands. Although the phylogenetic relationships among most of the taxa in clade I are still not well resolved, our use of more nuclear and plastid regions was helpful in recovering the African clade with higher bootstrap support than previously reported (Will and Claßen-Bockhoff, 2017).

#### Subgen. Heterosphace

This group comprises three supported subclades (supplementary 1). 1) Subclade I-A includes species from both northern and southern Africa. *Salvia nilotica* and *S. somalensis*, two species distributed in Tanzania and Ethiopia, respectively, were recovered as successive sisters to the southern African clade. 2) Subclade I-B comprises several species of *Salvia* from North America, formerly classified in section *Salviastrum* Scheele. Kriebel et al. (2019) argued that dispersal to eastern North America (sect. *Salviastrum* of subgen. “*Heterosphace*”) from the Eastern Hemisphere lineage occurred during the mid-Miocene. 3) The *S. verticillata* group: *Salvia taraxacifolia* was recovered as sister to the *S. verticillata* group (BS = 81%) consisting of *S. verticillata*, *S. judaica*, and *S. russellii*. The latter clade was sister (BS = 95%) to the southern African clade. Previous studies (Will and Claßen-Bockhoff, 2014; Will and Claßen-Bockhoff, 2017) failed to resolve the phylogenetic position of *S. taraxacifolia* (Mediterranean element) or with low support (kriebel eta l., 2019) within the *S. verticillata* group. Our results from the combined nuclear and plastid data recovered *S. taraxacifolia* as sister *to* the *S. verticillata* group with high support. *S. nilotica* placed as sister to the southern African clade species.

Most of the *Salvia* species in clade I (Europe, Madagascar, Central Asia), along with Iranian species of *Salvia*, are placed in subclades I-C and I-D and are classified as subgen. *Sclarea* and *Salvia* following recent treatments (Drew et al., 2017; Kriebel et al., 2019). Although phylogenetic relationships among species of *Salvia* in these subclades are mostly unresolved, our increased taxon sampling was helpful in determining the phylogenetic position of Iranian *Salvia* species and provides evolutionary insights and rationale for improving the taxonomy of *Salvia*.

#### Clade II

As noted above, clade II comprises species from North and South America, within which approximately half of all *Salvia* species are distributed. Because relationships among species from this geographic region were not the focus of this study, our sampling from this region is more limited than from central and western Asia, Europe, and Africa. Relationships are therefore largely unresolved. For more detail about relationships among species of *Salvia* from the Americas, we refer the reader to (Walker et al., 2015; Fragoso-Martínez 2018; Kriebel et al., 2019)

#### Clade III

Clade III encompasses species distributed in northern Africa and southwestern Asia. Within this clade, species are mostly dwarf shrubs with smaller flowers than those of other *Salvia* clades. In the Iran flora, most species are distributed in the southern region of Iran (25-27 N°). We present here the best-resolved phylogeny obtained for clade III with the most comprehensive taxon sampling to date; our nuclear data set fully resolved relationships with BS = 100%. *Salvia majdae*, formerly classified as the monotypic genus *Zhumeria*, is an endemic aromatic shrub in southern Iran. Based on a recent study (Soltanipour et al., 2020), *S. majdae* is reported as an endangered species on the IUCN Red List based on Extent of Occurrence (EO) and Area Of Occupancy (AOO). *Salvia aristata* is another endemic species of Iran in this clade placed as the sister species of *S. majdae*. This species has a different habit (herabaceous perennial) from the remaining species in clade III, as well as a larger distribution in Iran.

#### Clade IV

Clade IV is restricted to eastern Asia. Notably, *S. glutinosa*, which is distributed in northern Iran and western Europe, forms a clade with species from eastern Asia. *Salvia glutinosa* also shares similar traits with *S. nubicola* in corolla color (yellow with brown-purple spots on the lower lip) and with a clade of *S. koyamae*, *S. glabrescense*, and *S. nipponica* in leaf form and flower shape (Hu et al., 2018). Thus, *S. glutinosa* may have historically had a larger distribution than currently displayed. It is likely that *S. glutinosa* is a relict Arcto-Trertiary element and that the Euxine-Hyrcanian province (western Europe, northern Iran) was a refugium for this species (Browicz 1987, Akhani et al., 2010). Ecological niche modeling projecting into the past may enable a more complete view of the past distribution of *S. glutinosa*. A more detailed view of the phylogeny of *Salvia* from eastern Asia is found in (Hu et al., 2018; Hu et al., 2020xx.

### 4.2. Divergence times

Our estimate for the date of origin of *Salvia* (31 Ma, 95% HPD = 37.6-27.5 Ma) is consistent with previous studies (Drew et al., 2012; Drew et al., 2017), which is not surprising given that the calibration used here was based on Drew et al. (2012) from analysis on a larger taxonomic scale of Nepethoideae based on a fossil fruit of *Melissa* from the Early-Middle Oligocene.

The Qinghai-Tibetan Plateau (QTP) underwent four periods of uplift: 25-17 Ma, 15-13 Ma, 8- 7 Ma, and 3.5-1.6 Ma. The major radiation for *Salvia* in eastern Asia in clade IV is estimated at 8-10 Ma, which coincides with the QTP uplift in the late Miocene. Our estimate for the crown age of eastern Asian *Salvia* (∼12 Ma) is consistent with that of Drew et al. (2017), but is younger than that reported in another recent study (Hu et al., 2018) on eastern Asian *Salvia* with an estimated date of ∼17 Ma. This inconsistency might be because of different taxon sampling, placement of calibrations, or prior distribution of the calibration node among the studies. Our data suggest that the QTP uplift played an important role in local diversification of *Salvia*, as it has for other plant genera in eastern Asia (Yao et al., 2016; Malik et al., 2017; Hu et al., 2018).

The Arabia-Eurasian collision in the Oligocene-early Miocene led to the emergence of the Alborz and Zagros Mountains in the Middle Miocene (15-12 Ma) in the Iranian plateau (Manafzadeh et al., 2016). The main stage of crustal thickening from the collision was ∼25 Ma, and the uplift of the Iranian plateau took place ∼15-12 Ma, with further uplift ∼5 Ma (Djamali et al., 2012; Manafzadeh et al., 2016). The emergence of these mountains coincides with the age of the MRCA of Iranian *Salvia* species. Formation and uplift of mountains can play an important role in evolutionary diversification through providing heterogeneous niches and landscapes. Therefore, we postulate that the emergence and uplift of the Iranian Mountains during the last ∼12 Ma, along with subsequent aridification (Manafzadeh et al., 2016; Folk et al., 2020), provided new ecological opportunities and habitat for *Salvia* diversification in Iran.

### 4.3. Ancestral state reconstruction

#### 4.3.1. Corolla length

Across *Salvia* phylogeny, there were multiple shifts from a corolla length longer than 25 mm to shorter corollas within and among clades. The MRCA of subg. *Calosphace* had a corolla length less than 25 mm, but multiple shifts from short (∼4 mm) to long (∼45 mm) corollas occurred in subclades of *Calosphace*. Based on the current study and previous reports (Wester and Claßen-Bockhoff, 2011; Wester et al., 2020), most *Salvia* species with an average corolla length of 22.3±6.5 mm are visited by bees, while bird pollinators are more attracted to flowers with an average corolla length of 31±9.5 mm. Floral construction is associated with the type of pollinators in *Salvia* (Wester et al., 2020), and an overall correlation between flower size and pollinator is not expected across all *Salvia* lineages. For instance, *S. blepharochlaena*, which is distributed in Turkey, is melittophilous, but has a long corolla. *Salvia purpurea* in subgen. *Audibertia* has an intermediate flower (pollinated by bees and hummingbirds); *S. purpurea* has a long corolla (19-36 mm) and long flower tube that is characteristic of hummingbird-pollinated *Salvia* species, but the flower has the wide landing platform of a bee-pollinated flower (Wester and Claßen-Bockhoff, 2011). Special flower traits like a short flower tube cause a phenotypic trade-off and adaptation to birds and bees, but if a short tube is combined with a narrow corolla opening, this combination of floral traits can generalize to both pollinators (Ohashi et al., 2021). We do not imply that corolla length is the only trait involved in *Salvia*–pollinator interactions; other factors, such as flower shape, tube length, and color, may also be involved in pollinator attraction and adaptation (Landis et al., 2018; Wessinger et al., 2019; Kriebel et al., 2020).

#### 4.3.2. Lever mechanism

*Salvia* species with an active lever mechanism are characterized by modified stamens. The lever is formed by elongation of the connective tissue that widens and separates the two thecae from each other. Levers have evolved several times in parallel both within and between clades I and II (Drew and Sytsma, 2012). In this study, we inferred the ancestral state based on two models considering different rates of evolution. Based on the ER model (Equal Rates), the ancestral state for *Salvia* is equivocal, and two alternative hypotheses may explain the distribution of the lever mechanism across *Salvia*. First, the MRCA of *Salvia* may have had a non-active lever mechanism, and a lever evolved independently multiple times in separate lineages. Alternatively, the ancestor of *Salvia* may have had an active lever mechanism, and several losses and reversals took place throughout the clade. Additionally, the HiSSE analysis, with the preference of the irreversible model for lever mechanism diversification, suggests that changes from non-active to active lever is not plausible or at the very least evolutionarily difficult. Therefore, we argue that a *Salvia* ancestor with an active lever is more probable than a non-active lever.

#### 4.3.3. Habit

The ancestral habit in *Salvia* is reconstructed here as equivocal. Several shifts from woody to herbaceous occurred within the main clades. This ambiguity might be due to diverse clades that transition frequently between woody and herbaceous, making it difficult to infer the state of the MRCA of *Salvia*.

### 4.4. Corolla length evolution

#### 4.4.1. Disparity Through Time

The value obtained (MDI = 0.21) for the disparity of corolla length reflects a high rate of diversification and morphological lability in related species. The positive MDI value for disparity shows that most of the variation in corolla size is within subclades, while a negative MDI indicates higher disparity among subclades, which is traditionally interpreted as adaptive radiation (Harmon et al., 2010). Increasing disparity in corolla length during the last 10 M years of *Salvia* evolution coincides with a number of geological events, including the Andean uplift, Mexican vulcanization (clade II), the uplift of the QTP (clade IV), and the uplift of the Zagros Mountains (occupied by species in clade I; Ferrari et al., 2012; Yao et al., 2016; Manafzadeh et al., 2016), all major geological events that may have profoundly shaped *Salvia* evolution worldwide.

Our positive value of MDI and support for the OU model contrast with traditional interpretations of adaptive radiation in which MDI is negative through phenotypic diversification with the Early Burst (EB) model of diversification (Harmon et al., 2010). In addition, the DDD analyses do not support density-independent lineage diversification, and the Yule model was selected as the best model with no apparent slowdown in *Salvia* diversification. In the classic definition of adaptive radiation, the rate of diversification first increases due to access to new niche space, followed by slow diversification as niche space fills (Rabosky, 2013; Gillespie et al., 2020). However, Augilee et al. (2018) argued that ecological niche filling as an explanation for negative-dependent diversity should be treated with caution because biotic (competition) and abiotic factors (landscape dynamics) can correspond to species diversity in different stages of a clade’s history.

#### 4.4.2. Corolla length diversification

Floral traits have played a key role in enhancing angiosperm diversification (e.g., Stebbins, 1970; Fenster et al., 2014; Armbruster, 2014; Van der Neit and Johnson, 2014), and some floral characters (corolla length, corolla tube length, corolla shape, and flower color) are associated with pollinator interactions. The positive effects of certain floral traits on the effectiveness of one group of pollinators relative to others occurs most often in bilaterally symmetrical flowers (Ollerton, 2009; Armbruster, 2014; Wester et al., 2020). The rate of evolution of corolla length in one clade of *Calosphace* (clade II) was significantly higher than in other clades. Detection of correlated rate shifts in this clade implies that changes in corolla length may have enabled an adaptive radiation in this clade. Species in this clade are mostly distributed in South and Central America, including Bolivia, Mexico, Peru, and Argentina, and include hummingbird-pollinated species with several shifts to bee pollination (Fragoso- Martinez et al., 2017; Kriebel et al., 2019). Therefore, corolla size may be a putatively adaptive trait that facilitated pollinator-flower interactions in this clade.

### 4.5. Lever-mechanism-dependent diversification

The special lever-mechanism pollination system in *Salvia* has been hypothesized to have played a major role in *Salvia* diversification (Claßen-Bockhoff et al., 2004; Drew and Sytsma, 2012). The functionality and structure of the lever mechanism were tested through field investigation and biomechanical experiments (Claßen-Bockhoff et al., 2004; Wester and Claßen-Bockhoff, 2004; Reith et al., 2007; Drew and Sytsma, 2012; Zheng et al., 2015). However, the actual effect of the lever mechanism on diversification has not been previously investigated. We examined this hypothesis across our phylogeny using a Hidden Markov Model implemented in the HiSSE package. The best model fitted was the HiSSE model with irreversible transitions among states. The lever mechanism likely has an important impact on pollination success (Classen-Bockhoff et al., 2004; Zheng et al., 2015; Kriebel et al., 2019) and may have influenced diversification, but we did not find any evidence for a direct association of lever mechanism with *Salvia* diversification. Characters not measured here, including flower shape features that are associated with the observed state, were likely influential as well (Kriebel et al., 2020). Based on the HiSSE analysis, we suggest that emphasis on the lever mechanism alone as the key promotor of diversification in *Salvia* may be misplaced and that other phenotypic characters, especially other floral traits, should also be considered and examined across the phylogeny. We should take into account that there might be shortcomings and insufficient information in macroevolutionary models and that trees for extant species may not permit the precise reconstruction of historical diversification (Louca and Pennell, 2020). However, Helmsetter et al. (2021) argue that recent more complex models can provide additional information and overcome the problems of relying on time trees for extant species. An important issue for future studies in understanding *Salvia* evolutionary history is assessing the effect of other floral traits on diversification via the reconstruction of more robust phylogenetic trees using more genes and species.

## Acknowledgement

We are thankful to Matthew Gitzendanner and Evgeny Mavrodiev, Florida Museum of Natural History, for help with analyses and general lab assistance.

## References

1. Aguilée, R., Gascuel, F., & Lambert, A. (2018). abiotic controls of speciation and extinction rates. Nature Communications, 1–13. https://doi.org/10.1038/s41467-018-05419-7.

2. Akaike, H. 1974.Anewlook at the statistical model identification. IEEE Trans. Autom. Control 19:716–723.

3. Akhani, H., Djamali, M., Ghorbanalizadeh, A., & And, E. R. (2010). Plant biodiversity of Hyrcanian relict forests, N Iran: An overview of the flora, vegetation, palaeoecology and conservation. Pakistan Journal of Botany, 42: 231–258.

4. Alfaro, M.E., Brock, C.D., Banbury, B.L. et al. Does evolutionary innovation in pharyngeal jaws lead to rapid lineage diversification in labrid fishes?. BMC Evol Biol 9, 255 (2009). https://doi.org/10.1186/1471-2148-9-255

5. Armbruster, W. S. (2014). Floral specialization and angiosperm diversity: Phenotypic divergence, fitness trade-offs and realized pollination accuracy. AoB PLANTS, 6, 1–24. https://doi.org/10.1093/aobpla/plu003

6. Beaulieu, J. M., & O’Meara, B. C. (2016). Detecting hidden diversification shifts in models of trait-dependent speciation and extinction. Systematic Biology, 65(4), 583–601. https://doi.org/10.1093/sysbio/syw022

7. Bolòs, O. de 1924-2007. (1990). Flora manual dels països catalans. https://doi.org/10.1017/CBO9781107415324.004

8. Bouckaert, R., Heled, J., Kühnert, D., Vaughan, T., Wu, C.-H., Xie, D., Drummond, A. J. (2014). BEAST 2: A Software Platform for Bayesian Evolutionary Analysis. PLOS Computational Biology, 10(4), 1–6. https://doi.org/10.1371/journal.pcbi.1003537

9. Browicz, K. (1989). Chorology of the Euxinian and Hyrcanian element in the woody flora of Asia. Plant Systematics and Evolution, 162(1), 305–314. https://doi.org/10.1007/BF00936923

10. Caetano, D. S., O’Meara, B. C., & Beaulieu, J. M. (2018). Hidden state models improve state-dependent diversification approaches, including biogeographical models. Evolution, 72(11), 2308–2324. https://doi.org/10.1111/evo.13602

11. Celep, F., Atalay, Z., Dikmen, F., Doğan, M., & Classen-Bockhoff, R. (2014). Flies as pollinators of melittophilous *Salvia* species (Lamiaceae). American Journal of Botany, 101(12), 2148– 2159. https://doi.org/10.3732/ajb.1400422

12. Claßen-Bockhoff, R., Speck, T., Tweraser, E., Wester, P., Thimm, S., & Reith, M. (2004). The staminal lever mechanism in *Salvia* L. (Lamiaceae): A key innovation for adaptive radiation? Organisms Diversity and Evolution, 4(3), 189–205. https://doi.org/10.1016/j.ode.2004.01.004

13. Claβen-Bockhoff, R., Wester, P., & Tweraser, E. (2003). The Staminal Lever Mechanism in *Salvia* L. (Lamiaceae) - a Review. Plant Biology, 5(1), 33–41. https://doi.org/10.1055/s-2003-37973

14. Crane, P. R., Friis, E. M., & Pedersen, K. R. (1995). The origin and early diversification of angiosperms. Nature, 374(6517), 27–33.

15. Crepet, W. L. (2000). Progress in understanding angiosperm history, success, and relationships: Darwin&#039;s abominably “perplexing phenomenon.” Proceedings of the National Academy of Sciences, 97(24), 12939 LP-12941. Retrieved from http://www.pnas.org/content/97/24/12939.abstract

16. Deng, T., Nie, Z., Drew, B. T., Volis, S., & Kim, C. (2015). Does the Arcto-Tertiary Biogeographic Hypothesis Explain the Disjunct Distribution of Northern Hemisphere Herbaceous Plants ? The Case of Meehania ( Lamiaceae ), 1–18. https://doi.org/10.1371/journal.pone.0117171

17. Djamali, M., Baumel, A., Brewer, S., Jackson, S. T., Kadereit, J. W., López-vinyallonga, S., Simakova, A. (2012). Review of Palaeobotany and Palynology Ecological implications of Cousinia Cass . ( Asteraceae ) persistence through the last two glacial – interglacial cycles in the continental Middle East for the Irano-Turanian fl ora. Review of Palaeobotany and Palynology, 172, 10–20. https://doi.org/10.1016/j.revpalbo.2012.01.005

18. Dodd, M. E., Silvertown, J., Chase, M. W. (1999). Analysis of trait evolution and species diversity variation among angiosperm families. Evolution, 53(3), 732–744.

19. Drew, B. T., & Sytsma, K. J. (2012). Phylogenetics, biogeography, and staminal evolution in the tribe Mentheae (Lamiaceae). American Journal of Botany, 99(5), 933–953. https://doi.org/10.3732/ajb.1100549

20. Drew, B. T., González-gallegos, J. G., Xiang, C., Kriebel, R., Drummond, C. P., Walker, J. B., & Sytsma, K. J. (2017). *Salvia* united : The greatest good for the greatest number. TAXON, 66(1), 133–145.

21. Drummond, A. J., & Rambaut, A. (2007). BEAST : Bayesian evolutionary analysis by sampling trees, 8, 1–8. https://doi.org/10.1186/1471-2148-7-214

22. Epling, C. (1938-1939) A revision of *Salvia*, subgenus *Calosphace*. Feddes Rep. Beith., 110, 1–383.

23. Etienne, R. S., & Haegeman, B. (2012). A conceptual and statistical framework for adaptive radiations with a key role for diversity dependence. American Naturalist, 180(4). https://doi.org/10.1086/667574

24. Fenster, C. B., Armbruster, W. S., Wilson, P., Dudash, M. R., & Thomson, J. D. (2004). Pollination syndromes and floral specialization. Annual Review of Ecology, Evolution, and Systematics, 35, 375–403.https://doi.org/10.1146/annurev.ecolsys.34.011802.132347

25. Fernández-Mazuecos, M., Blanco-Pastor, J. L., Gómez, J. M., & Vargas, P. (2013). Corolla morphology influences diversification rates in bifid toadflaxes (Linaria sect. Versicolores). Annals of Botany, 112(9), 1705–1722. https://doi.org/10.1093/aob/mct214

26. Ferrari, L., Orozco-Esquivel, T., Manea, V., & Manea, M. (2012). The dynamic history of the Trans-Mexican Volcanic Belt and the Mexico subduction zone. Tectonophysics, 522–523, 122–149. https://doi.org/10.1016/j.tecto.2011.09.018

27. FitzJohn, R. G. (2012). Diversitree : comparative phylogenetic analyses of diversification in R. Methods in Ecology and Evolution, (3), 1084–1092. https://doi.org/10.1111/j.2041-210X.2012.00234.x

28. FitzJohn, R. G., Pennell, M. W., Zanne, A. E., Stevens, P. F., Tank, D. C., & Cornwell, W. K. (2014). How much of the world is woody ?, (Theophrastus 1916), 1266–1272. https://doi.org/10.5061/dryad.63q27/2

29. Fragoso-Martínez, I., Martínez-Gordillo, M., Salazar, G. A., Sazatornil, F., Jenks, A. A., García Peña, M. del R., Granados Mendoza, C. (2018). Phylogeny of the Neotropical sages (*Salvia* subg. Calosphace; Lamiaceae) and insights into pollinator and area shifts. Plant Systematics and Evolution, 304(1), 43–55. https://doi.org/10.1007/s00606-017-1445-4

30. Folk, R. A., C. M. Siniscalchi, D. E. Soltis. 2020. Angiosperms at the edge: Extremity and diversity. Plant, Cell & Environment 43: 2871– 2893. https://doi.org/10.1111/pce.13887

31. Gillespie, R. G., Bennett, G. M., De Meester, L., Feder, J. L., Fleischer, R. C., Harmon, L. J Wogan, G. O. U. (2020). Comparing Adaptive Radiations Across Space, Time, and Taxa. Journal of Heredity, 111(1), 1–20. https://doi.org/10.1093/jhered/esz064

32. Gómez, J. M., Torices, R., Lorite, J., Klingenberg, C. P., & Perfectti, F. (2016). The role of pollinators in the evolution of corolla shape variation, disparity and integration in a highly diversified plant family with a conserved floral bauplan. Annals of Botany, 117(5), 889–904. https://doi.org/10.1093/aob/mcv194

33. Givnish, T. J. (2015). Adaptive radiation versus “ radiation ” and “ explosive diversification ”: why conceptual distinctions are fundamental to understanding evolution, New Phytologist, 207, 297–303..

34. Han, T.-S., Zheng, Q.-J., Onstein, R. E., Rojas-Andrés, B. M., Hauenschild, F., Muellner-Riehl, A. N., & Xing, Y.-W. (2020). Polyploidy promotes species diversification of Allium through ecological shifts. New Phytologist, 225(1), 571–583. https://doi.org/10.1111/nph.16098

35. Hansen, T. F. (1997). Stabilizing selection and the comparative analysis of adaptation. Evolution, 51(5), 1341–1351. https://doi.org/10.1111/j.1558-5646.1997.tb01457.x.

36. Harder, L. D., & Johnson, S. D. (2009). Darwin’ s beautiful contrivances : evolutionary and functional evidence. New Phytologist, 183, 530–545. https://doi.org/10.1111/j.1469-8137.2009.02914.x

37. Harley, R. M., Atkins, S., Budantsev, A. L., Cantino, P. D., Conn, B. J., Grayer, R., Upson, T. (2004). Labiatae. In J. W. Kadereit (Ed.), Flowering Plants·Dicotyledons: Lamiales (except Acanthaceae including Avicenniaceae) (pp. 167–275). Berlin, Heidelberg: Springer Berlin Heidelberg. https://doi.org/10.1007/978-3-642-18617-2_11

38. Harmon, L. J., Schulte, J. A., Larson, A., & Losos, J. B. (2003). Tempo and Mode of Evolutionary Radiation in Iguanian Lizards. Science, 301(5635), 961–964. https://doi.org/10.1126/science.1084786

39. Harmon Luke J, Jason T Weir, Chad D Brock, Richard E Glor, and Wendell Challenger. 2008. GEIGER: investigating evolutionary radiations. Bioinformatics 24, 129–131.

40. Harmon, L. J., Losos, J. B., Jonathan Davies, T., Gillespie, R. G., Gittleman, J. L., Bryan Jennings, W., Mooers, A. T. (2010). Early bursts of body size and shape evolution are rare in comparative data. Evolution, 64(8), 2385–2396. https://doi.org/10.1111/j.1558-5646.2010.01025.x

41. Harrington, S., & Reeder, T. W. (2017). Rate heterogeneity across Squamata, misleading ancestral state reconstruction and the importance of proper null model specification, 30, 313–325. https://doi.org/10.1111/jeb.13004

42. Hedge, I.C. (1982a). Salvia L. In: Davis PH (ed.) Flora of Turkey and the East Aegean Islands, 7, 400–461. Edinburgh Univ. Press.

43. Hedge, I.C. (1982b). Salvia L. In: Rechinger KH (ed.), Flora Iranica 150, 403–476, Akademische Druck und Verlagsanstalt, Graz.

44. Helmstetter, A. J., Glemin, S., Käfer, J., Zenil-Ferguson, R., Sauquet, H., de Boer, H. Condamine, F. L. (2021). Pulled Diversification Rates, Lineages-Through-Time Plots and Modern Macroevolutionary Modelling. BioRxiv. https://doi.org/10.1101/2021.01.04.424672

45. Hernández-Hernández, T., & Wiens, J. J. (2020). Why are there so many flowering plants? A multiscale analysis of plant diversification. American Naturalist, 195(6), 948–963. https://doi.org/10.1086/708273

46. Hu, G.-X., Takano, A., Drew, B. T., Liu, E.-D., Soltis, D. E., Soltis, P. S., Xiang, C.-L. (2018). Phylogeny and staminal evolution of *Salvia* (Lamiaceae, Nepetoideae) in East Asia. Annals of Botany, 122(4), 649–668. https://doi.org/10.1093/aob/mcy104

47. Hu, G.-X., Liu, E.-D., Wu, Z.-K., Sytsma, K. J., Drew, B. T., & Xiang, C.-L. (2020). Integrating DNA Sequences with Morphological Analysis Clarifies Phylogenetic Position of Salvia grandifolia (Lamiaceae): An Enigmatic Species Endemic to Southwestern China. International Journal of Plant Sciences, 181(8), 787–799. https://doi.org/10.1086/709134

48. Kaczorowski, R. L., Seliger, A. R., Gaskett, A. C., Wigsten, S. K., & Raguso, R. A. (2012). Corolla shape vs. size in flower choice by a nocturnal hawkmoth pollinator. Functional Ecology, 26(3), 577–587. https://doi.org/10.1111/j.1365-2435.2012.01982.x

49. Koutroumpa, K., Warren, B. H., Theodoridis, S., Coiro, M., Romeiras, M. M., Jiménez, A., & Conti, E. (2021). Geo-Climatic Changes and Apomixis as Major Drivers of Diversification in the Mediterranean Sea Lavenders (Limonium Mill.). Frontiers in Plant Science .https://doi.org/10.3389/fpls.2020.612258

50. Kay, K. M., & Sargent, R. D. (2009). The Role of Animal Pollination in Plant Speciation: Integrating Ecology, Geography, and Genetics. *Annual Review of Ecology*, Evolution, and Systematics, 40(1), 637–656. https://doi.org/10.1146/annurev.ecolsys.110308.120310

51. Kriebel, R., Drew, B. T., Drummond, C. P., González-Gallegos, J. G., Celep, F., Mahdjoub, M. M., Sytsma, K. J. (2019). Tracking temporal shifts in area, biomes, and pollinators in the radiation of *Salvia* (sages) across continents: leveraging anchored hybrid enrichment and targeted sequence data. American Journal of Botany, 106(4), 573–597. https://doi.org/10.1002/ajb2.1268

52. Kriebel, R., Drew, B., González-Gallegos, J. G., Celep, F., Heeg, L., Mahdjoub, M. M., & Sytsma, K. J. (2020). Pollinator shifts, contingent evolution, and evolutionary constraint drive floral disparity in *Salvia* (Lamiaceae): Evidence from morphometrics and phylogenetic comparative methods. Evolution, 74(7), 1335–1355. https://doi.org/10.1111/evo.14030

53. Lagomarsino, L. P., Condamine, F. L., Antonelli, A., Mulch, A., & Davis, C. C. (2016). The abiotic and biotic drivers of rapid diversification in Andean bellflowers (Campanulaceae). New Phytologist, 210(4), 1430–1442. https://doi.org/10.1111/nph.13920

54. Landis, J. B., Bell, C. D., Hernandez, M., Zenil-Ferguson, R., McCarthy, E. W., Soltis, D. E., & Soltis, P. S. (2018). Evolution of floral traits and impact of reproductive mode on diversification in the phlox family (Polemoniaceae). Molecular Phylogenetics and Evolution, 127(June), 878–890. https://doi.org/10.1016/j.ympev.2018.06.035

55. Landis, J. B., Toole, R. D. O., Ventura, K. L., & Gitzendanner, M. A. (2016). The Phenotypic and Genetic Underpinnings of Flower Size in Polemoniaceae The Phenotypic and Genetic Underpinnings of Flower Size in Polemoniaceae, 6(November), 1–20. https://doi.org/10.3389/fpls.2015.01144

56. Li, B., Cantino, P. D., Olmstead, R. G., Bramley, G. L. C., Xiang, C. L., Ma, Z. H., Zhang, D. X. (2016). A large-scale chloroplast phylogeny of the Lamiaceae sheds new light on its subfamilial classification. Scientific Reports, 6(October). https://doi.org/10.1038/srep34343

57. Li, P., Qi, Z. C., Liu, L. X., Ohi-Toma, T., Lee, J., Hsieh, T. H., Qiu, Y. X. (2017). Molecular phylogenetics and biogeography of the mint tribe Elsholtzieae (Nepetoideae, Lamiaceae), with an emphasis on its diversification in East Asia. Scientific Reports, 7(1), 1–12. https://doi.org/10.1038/s41598-017-02157-6

59. Louca, S., & Pennell, M. W. (2020). Extant timetrees are consistent with a myriad of diversification histories. Nature, 580(7804), 502–505.

60. Maddison, W. P., Midford, P. E., & Otto, S. P. (2007). Estimating a Binary Character’s Effect on Speciation and Extinction. Systematic Biology, 56(5), 701–710. Retrieved from http://dx.doi.org/10.1080/10635150701607033

61. Malik, S., Vitales, D., Qasim Hayat, M., Korobkov, A. A., Garnatje, T., & Vallès, J. (2017). Phylogeny and biogeography of Artemisia subgenus seriphidium (Asteraceae: Anthemideae). Taxon, 66(4), 934–952. https://doi.org/10.12705/664.8

62. Manafzadeh, S., Salvo, G., & Conti, E. (2013). A tale of migrations from east to west : the Irano-Turanian floristic region as a source of Mediterranean xerophytes, 1–14. https://doi.org/10.1111/jbi.12185.

63. Manafzadeh, S., Staedler, Y. M., & Conti, E. (2016). Visions of the past and dreams of the future in the Orient : the Irano-Turanian region from classical botany to evolutionary studies, 9. https://doi.org/10.1111/brv.12287

64. McGuire, J. A., Witt, C. C., Remsen, J. V., Corl, A., Rabosky, D. L., Altshuler, D. L., & Dudley, R. (2014). Molecular phylogenetics and the diversification of hummingbirds. Current Biology, 24(8), 910–916. https://doi.org/10.1016/j.cub.2014.03.016

65. Ogutcen, E., Hamper, B., & Vamosi, J. C. (2014). Diversification in monkeyflowers: an investigation of the effects of elevation and floral color in the genus Mimulus. International Journal of Evolutionary Biology, 2014, 382453. https://doi.org/10.1155/2014/382453

66. Ohashi, K., Jürgens, A., & Thomson, J. D. (2021). Trade-off mitigation : a conceptual framework for understanding fl oral adaptation in multispecies interactions, 6. https://doi.org/10.1111/brv.12754

67. Onstein, R. E. (2019). Darwin’s second “abominable mystery”: trait flexibility as the innovation leading to angiosperm diversity. New Phytologist. https://doi.org/10.1111/nph.16294

68. Paradis, E., Claude, J., & Strimmer, K. (2004). APE: Analyses of phylogenetics and evolution in R language. Bioinformatics, 20(2), 289–290. https://doi.org/10.1093/bioinformatics/btg412

69. Pavlopoulos, G. A., Hooper, S. D., Sifrim, A., Schneider, R., & Aerts, J. (2011). Medusa: A tool for exploring and clustering biological networks. BMC Research Notes, 4(1), 384. https://doi.org/10.1186/1756-0500-4-384

70. Phylogenomic Mining of the Mints Reveals Multiple Mechanisms Contributing to the Evolution of Chemical Diversity in Lamiaceae. (2018). Molecular Plant, 11(8), 1084– 1096. https://doi.org/10.1016/j.molp.2018.06.002

71. Pincheira-Donoso, D., Harvey, L. P., & Ruta, M. (2015). What defines an adaptive radiation? Macroevolutionary diversification dynamics of an exceptionally species-rich continental lizard radiation. BMC Evolutionary Biology, 15(1), 153. https://doi.org/10.1186/s12862-015-0435-9

72. Pobedimova, E.G. (1954). Salvia L. - In: Schischkin, B.K. (ed.), Flora of the USSR 21, 178–260. [ translated from Russian] Israel Prog. Sci. Transel., Jerusalem.

73. Pyron, R. A., & Burbrink, F. T. (2013). Phylogenetic estimates of speciation and extinction rates for testing ecological and evolutionary hypotheses. Trends in Ecology & Evolution, 28(12), 729–736. https://doi.org/10.1016/j.tree.2013.09.007

74. Rabosky, D. L. (2006). LASER: A Maximum Likelihood Toolkit for Detecting Temporal Shifts in Diversification Rates from Molecular Phylogenies. Evolutionary Bioinformatics, 2, 247–250. https://doi.org/10.1177/117693430600200024.

75. Rabosky, D. L. (2017). Phylogenetic tests for evolutionary innovation: The problematic link between key innovations and exceptional diversification. Philosophical Transactions of the Royal Society B: Biological Sciences, 372(1735). https://doi.org/10.1098/rstb.2016.0417

76. Rabosky, D. L., & Goldberg, E. E. (2017). FiSSE: A simple nonparametric test for the effects of a binary character on lineage diversification rates. Evolution, 71(6), 1432–1442. https://doi.org/10.1111/evo.13227

77. Rabosky, D. L., & Goldberg, E. E. (2015). Model Inadequacy and Mistaken Inferences of Trait- Dependent Speciation. Systematic Biology, 64(2), 340–355. https://doi.org/10.1093/sysbio/syu131

78. Rabosky, D. L., Grundler, M., Anderson, C., Title, P., Shi, J. J., Brown, J. W., Larson, J. G. (2014). BAMMtools : an R package for the analysis of evolutionary dynamics on phylogenetic trees, 701–707. https://doi.org/10.1111/2041-210X.12199

79. Rabosky, Daniel L. 2014. “Automatic Detection of Key Innovations, Rate Shifts, and Diversity- Dependence on Phylogenetic Trees.” PLoS ONE 9(2).

80. Reith, M., Baumann, G., Claßen-Bockhoff, R., & Speck, T. (2007). New insights into the functional morphology of the lever mechanism of *Salvia pratensis* (Lamiaceae). Annals of Botany, 100(2), 393–400. https://doi.org/10.1093/aob/mcm031

81. Revell, L. J. (2012). phytools : an R package for phylogenetic comparative biology ( and other things ), Methods in Ecology and Evolution, 3: 217–223. https://doi.org/10.1111/j.2041-210X.2011.00169.x

82. Ricklefs, R. E. (2007). Estimating diversification rates from phylogenetic information. Trends in Ecology and Evolution, 22(11), 601–610. https://doi.org/10.1016/j.tree.2007.06.013

83. Sauquet, H., Balthazar, M. Von, Magallón, S., Doyle, J. A., Endress, P. K., Bailes, E. J., … Schönenberger, J. (2017). The ancestral flower of angiosperms and its early diversification. Nature Communications, 8. https://doi.org/10.1038/ncomms16047

84. Seelanan, T., Schnabel, A., & Wendel, J. F. (1997). Congruence and Consensus in the Cotton Tribe (Malvaceae). Systematic Botany, 22(2), 259–290. https://doi.org/10.2307/2419457

85. Serrano-Serrano, M. L., Perret, M., Guignard, M., Chautems, A., Silvestro, D., & Salamin, N. (2015). Decoupled evolution of floral traits and climatic preferences in a clade of Neotropical Gesneriaceae. BMC Evolutionary Biology, 15(1), 1–12. https://doi.org/10.1186/s12862-015-0527-6

86. Smith, S. A., & O’Meara, B. C. (2012). treePL: divergence time estimation using penalized likelihood for large phylogenies. Bioinformatics, 28(20), 2689–2690. Retrieved from http://dx.doi.org/10.1093/bioinformatics/bts492

87. Smith, S. D. (2010). Using phylogenetics to detect pollinator-mediated floral evolution. New Phytologist, 354–363.

88. Smith, S. D., & Kriebel, R. (2018). Convergent evolution of floral shape tied to pollinator shifts in *Iochrominae* (Solanaceae)*. Evolution, 72(3), 688–697. https://doi.org/10.1111/evo.13416.

89. Soltanipour, M.A, Jamzad, Z., Jalili, A., Mahmoodi, M (2020). The conservation status of *Zhumeria majdae*, an endemic species of Iran. Iran Nature. https://dx.doi.org/10.22092/irn.2021.121273

90. Soltis, D. E., Bell, C. D., Kim, S., & Soltis, P. S. (2008). Origin and Early Evolution of Angiosperms. Annals OF THe New York A Cademy of Sciences, 25, 3–25. https://doi.org/10.1196/annals.1438.005

91. Soltis, P.S. & D.E. Soltis. 2004. Phylogeny and evolution of the angiosperms. Am. J. Botany 91: 1614–1626.

92. Soltis, D. E., Mort, M. E., Latvis, M., Mavrodiev, E. V., O’Meara, B. C., Soltis, P. S., Burleigh, J. G., & Rubio de Casas, R. (2013). Phylogenetic relationships and character evolution analysis of Saxifragales using a supermatrix approach. American journal of botany, 100(5), 916–929. https://doi.org/10.3732/ajb.1300044

93. Soltis PS, Soltis DE. 2014.Flower diversity and angiosperm diversification. In:Riechmann JL, Wellmer F, eds.Flower development: methods and protocols. NewYork, NY, USA: Springer, 85–102.

94. Stebbins, G. L. (1970). Adaptive radiation of reproductive characteristics in angiosperms, 4012 i: pollination mechanisms. Annual Review of Ecology and Systematics, 1, 307–326.

95. Takano, A. (2017). Taxonomic study on Japanese Salvia (Lamiaceae): Phylogenetic position of S. akiensis, and polyphyletic nature of *S. lutescens* var. intermedia. PhytoKeys, 80, 87–104. https://doi.org/10.3897/phytokeys.80.11611

96. Takano, A., & Okada, H. (2011). Phylogenetic relationships among subgenera , species , and varieties of Japanese *Salvia* L . ( Lamiaceae ), 245–252. https://doi.org/10.1007/s10265-010-0367-9

97. The Angiosperm Phylogeny Group III. (2009). An update of the Angiosperm Phylogeny Group Classification for the orders and families of flowering plants: APG III. Botanical Journal of the Linnean Society, 161(2), 105–121. https://doi.org/10.1111/j.1095-8339.2009.00996.x

98. Vamosi, J. C., Vamosi, S. M., Van der Niet, T., & Johnson, S. D. (2011). Factors influencing diversification in angiosperms: At the crossroads of intrinsic and extrinsic traits. American Journal of Botany, 98(3), 460–471. https://doi.org/10.3732/ajb.1000311

99. Van der Niet, T., & Johnson, S. D. (2012). Phylogenetic evidence for pollinator-driven diversification of angiosperms. Trends in Ecology and Evolution, 27(6), 353– 361.https://doi.org/10.1016/j.tree.2012.02.002

100. Van Der Niet, T., Peakall, R., & Johnson, S. D. (2014). Pollinator-driven ecological speciation in plants: New evidence and future perspectives. Annals of Botany, 113(2), 199–211. https://doi.org/10.1093/aob/mct290

101. Wagstaff, S. J., Olmstead, R. G., & Cantino, P. D. (1995). Parsimony analysis of cpDNA restriction site variation in subfamily Nepetoideae (Labiatae). American Journal of Botany, 82(7), 886–892. https://doi.org/10.1002/j.1537-2197.1995.tb15705.x

102. Wagstaff, S. J., Olmstead, R. G., & Cantino, P. D. (1995). Parsimony analysis of cpDNA restriction site variation in subfamily Nepetoideae (Labiatae). American Journal of Botany, 82(7), 886–892. https://doi.org/10.1002/j.1537-2197.1995.tb15705.x

103. Walker, J. B., & Sytsma, K. J. (2007). Staminal evolution in the genus *Salvia* (Lamiaceae): Molecular phylogenetic evidence for multiple origins of the staminal lever. Annals of Botany, 100(2), 375–391. https://doi.org/10.1093/aob/mcl176

104. Walker, J. B., Sytsma, K. J., Treutlein, J., & Wink, M. (2004). *Salvia* (Lamiaceae) is not monophyletic: Implications for the systematics, radiation, and ecological specializations of *Salvia* and tribe Mentheae. American Journal of Botany, 91(7), 1115–1125. https://doi.org/10.3732/ajb.91.7.1115

105. Walker, J. B., Drew, B. T., & Sytsma, K. J. (2015). Unravelling Species Relationships and Diversification within the Iconic California Floristic Province Sages (Salvia Subgenus Audibertia, Lamiaceae). Systematic Botany, 40(3), 826–844. https://doi.org/10.1600/036364415X689285

106. Wester, P., & Claßen-Bockhoff, R. (2007). Floral diversity and pollen transfer mechanisms in bird-pollinated *Salvia* species. Annals of Botany, 100(2), 401–421. https://doi.org/10.1093/aob/mcm036

107. Wester, P., & Claßen-Bockhoff, R. (2011). Pollination Syndromes of New World *Salvia* Species with Special Reference to Bird Pollination ^1^. Annals of the Missouri Botanical Garden, 98(1), 101–155. https://doi.org/10.3417/2007035

108. Wester, P., Cairampoma, L., Haag, S., Schramme, J., Neumeyer, C., & Claßen-Bockhoff, R. (2020). Bee exclusion in bird-pollinated salvia flowers: The role of flower color versus flower construction. International Journal of Plant Sciences, 181(8), 770–786. https://doi.org/10.1086/709132

109. Wessinger, C. A., Rausher, M. D., & Hileman, L. C. (2019). Adaptation to hummingbird pollination is associated with reduced diversification in PenstemLandis, J. B., Bell, Charles D.Wessinger, C. A., Rausher, M. D., & Hileman, L. C. (2019). Adaptation to hummingbird pollination is associated with reduced diversifi. Evolution Letters, 3(5), 521–533. https://doi.org/10.1002/evl3.130

110. Will, M., & Claßen-Bockhoff, R. (2014). Why Africa matters: Evolution of Old World *Salvia* (Lamiaceae) in Africa. Annals of Botany, 114(1), 61–83. https://doi.org/10.1093/aob/mcu081

111. Will, M., & Claßen-Bockhoff, R. (2017). Time to split *Salvia* s.l. (Lamiaceae) – New insights from Old World Salvia phylogeny. Molecular Phylogenetics and Evolution, 109, 33–58. https://doi.org/10.1016/j.ympev.2016.12.041

112. Will, M., Schmalz, N., & Classen-bockhoff, R. (2015). Towards a new classification of Salvia s. l.: (re) establishing the genus Pleudia Raf ., 1–15. https://doi.org/10.3906/bot-1405-34

113. Wu, H., Ma, P.-F., Li, H.-T., Hu, G.-X., & Li, D.-Z. (2021). Comparative plastomic analysis and insights into the phylogeny of *Salvia* (Lamiaceae). Plant Diversity, 43(1), 15–26. https://doi.org/10.1016/j.pld.2020.07.004

114. Yao, G., Drew, B. T., Yi, T. S., Yan, H. F., Yuan, Y. M., & Ge, X. J. (2016). Phylogenetic relationships, character evolution and biogeographic diversification of *Pogostemon* s.l. (Lamiaceae). Molecular Phylogenetics and Evolution, 98, 184–200. https://doi.org/10.1016/j.ympev.2016.01.02

115. Zanne, A. E. D. C. Tank, W. K. Cornwell, J. M. Eastman, S. A. Smith, R. G. FitzJohn, D. J. McGlinn, B. C. O’Meara, A. T. Moles, P. B. Reich, D. L. Royer, D. E. Soltis, P. F. Stevens, M. Westoby, I. J. Wright, L. Aarssen, R. I. Bertin, A. Calaminus, R. Govaerts, F. Hemmings, M. R. Leishman, J. Oleksyn, P. S. Soltis, N. G. Swenson, L. Warman, J. M. Beaulieu. 2014. Into the cold - three keys to radiation of angiosperms into freezing environments. Nature doi:10.1038/nature12872

116. Zhang, B., Claßen-Bockhoff, R., Zhang, Z. Q., Sun, S., Luo, Y. J., & Li, Q. J. (2011). Functional implications of the staminal lever mechanism in *Salvia cyclostegia* (Lamiaceae). Annals of Botany, 107(4), 621–628. https://doi.org/10.1093/aob/mcr011

