## Supplementary figures and images for "Towards a global perspective for *Salvia* L: Phylogeny, diversification, and floral evolution"

### supplemental Figure 1

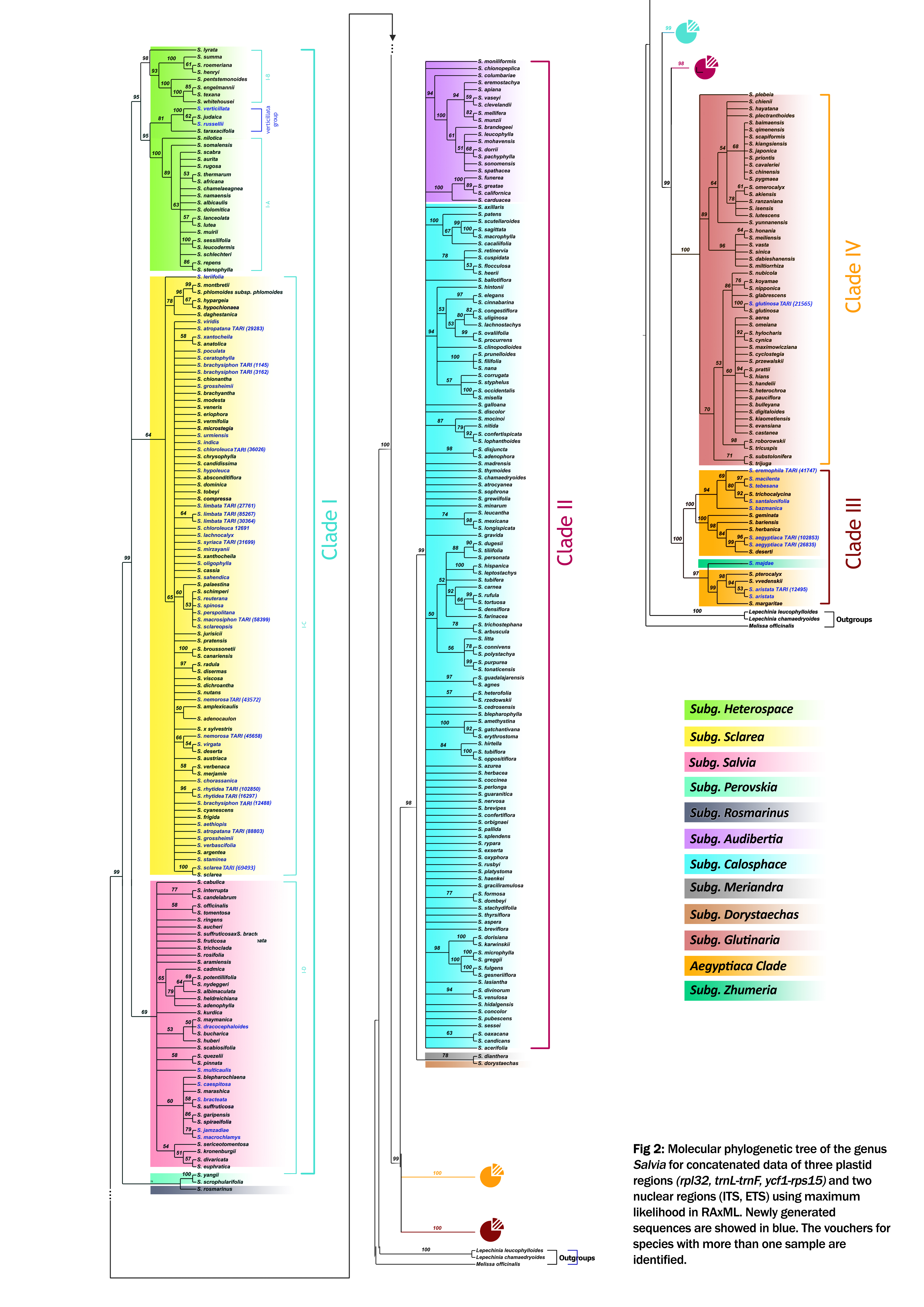
